# Aquaporin-4 and Caveolin-1 as Mediators of Fibrinogen-Driven Cerebrovascular Pathology in Cerebral Amyloid Angiopathy

**DOI:** 10.1101/2024.11.11.623066

**Authors:** Vishal Singh, Nicholas Rochakim, Francesca Ferraresso, Khushi Garg, Arnab Choudhury, Christian J. Kastrup, Hyung Jin Ahn

**Author notes:** Correspondence. Hyung Jin Ahn **Email:**.

## Abstract

Cerebral Amyloid Angiopathy (CAA), characterized by amyloid-β (Aβ) accumulation within perivascular spaces (PVS), contributes to vascular damage and inflammation in Alzheimer’s disease (AD). Despite its significance, the mechanisms driving Aβ deposition in PVS and the resulting vascular pathology remain poorly understood. Growing evidence suggests that fibrinogen, the main component in blood clots, interacts with Aβ and exacerbates inflammation in AD. Fibrinogen also co-deposits with Aβ in the PVS of CAA-positive vessels in the brains of hereditary CAA patients. However, the mechanisms by which fibrinogen contributes to cerebrovascular impairment remain poorly understood. To investigate this, we used TgSwDI transgenic mice, which develop robust CAA pathology, and observed a significant increase in fibrin(ogen) extravasation and colocalization with Aβ in the PVS. Moreover, we observed a significant aquaporin-4 (AQP4) depolarization in CAA-laden blood vessels of TgSwDI mice, which correlated with fibrin(ogen)-Aβ colocalization. Given AQP4 crucial role in Aβ clearance through glymphatic pathway, its depolarization may disrupt critical Aβ clearance, thereby exacerbating CAA pathology. Additionally, Caveolin-1, a protein involved in non-specific transcytosis across the endothelium, significantly increased with age in TgSwDI mice and correlated with fibrin(ogen) extravasation. To further explore the relationship between fibrin(ogen) and these cerebrovascular alterations, we depleted fibrinogen in TgSwDI mice using siRNA approach. This intervention resulted in decreased CAA, restored polarized expression of AQP4, reduced caveolin-1 levels, attenuated microglial activation, and improved spatial memory in fibrinogen-depleted TgSwDI mice. These findings suggest that targeting fibrinogen could be a promising strategy for mitigating CAA pathology and its associated cerebrovascular pathology.

**Significance Statement:** Our study uncovers the mechanism by which fibrin(ogen)-Aβ colocalization exacerbates CAA pathology. Our findings highlight the potential link between fibrinogen/ fibrin(ogen)-Aβ colocalization and AQP4 depolarization thereby exacerbating CAA pathology. The age-dependent increase of endothelial caveolin-1 could facilitate fibrin(ogen) extravasation, assisting the later to binds to Aβ in the perivascular space which ultimately induce microglial neuroinflammation and AQP4 depolarization, thus exacerbating CAA pathology. Furthermore, fibrinogen depletion could mitigate CAA severity, reduce microglial activation, restore AQP4 polarization and memory impairment. These results suggest that targeting fibrinogen and caveolin-1-mediated transcytosis may offer new strategies to address CAA-associated cerebrovascular pathology.

## Introduction

Brain vascular changes have been reported as an early pathological event in Alzheimer’s disease (AD) progression (1–4). Among these changes, cerebral amyloid angiopathy (CAA), characterized by the deposition of β-amyloid (Aβ) around cerebral blood vessels, plays a crucial role in vascular dysfunction in AD and is observed in over 80% of AD patients (5, 6). CAA contributes to a variety of vascular abnormalities, including altered cerebral blood flow, CAA-related inflammation, cerebral hemorrhages, and microinfarcts (7–10), all of which are linked to cognitive impairment in AD (11). Despite its high prevalence and clinical significance, the mechanisms driving CAA development and its contribution to neurodegeneration remain poorly understood. In particular, it is unclear how Aβ accumulates in perivascular spaces (PVS) and what downstream events lead to cerebrovascular damage.

Emerging evidence suggests that impaired clearance of Aβ through the glymphatic system and/or intramural periarterial drainage (IPAD) pathways contributes to its accumulation in CAA (12). A key component of glymphatic Aβ clearance is aquaporin-4 (AQP4), a main water channel protein highly enriched in astrocytic endfeet that facilitates cerebrospinal fluid (CSF)– interstitial fluid (ISF) exchange along perivascular routes (13–17). Disruption of AQP4 function has been implicated in reduced Aβ clearance and increased vascular deposition. Altered expression and depolarization of AQP4 have been observed in both AD and CAA patients (18–21). In mouse models of AD, loss of perivascular AQP4 impairs glymphatic transport and accelerates Aβ plaque formation (22). Similarly, human AD brains exhibit diminished perivascular AQP4 localization relative to cognitively normal individuals, leading to impaired glymphatic exchange and early Aβ accumulation (22). However, the mechanisms responsible for this loss of AQP4 polarity in AD and CAA remain to be elucidated.

A growing body of evidence has identified fibrinogen, a primary blood-clotting protein, as a potential contributor to neurovascular damage, neuroinflammation, and neuronal degeneration in both AD and CAA (23–26). Fibrinogen has been shown to bind Aβ, enhancing its aggregation and persistence, while also promoting neuroinflammatory responses. Notably, fibrinogen co-deposits with Aβ in the PVS of CAA-positive vessels in the brain of patients with hereditary forms of CAA (HCAA) (27). Moreover, fibrinogen extravasation has been associated with AQP4 loss and dementia in the brains of Idiopathic normal pressure hydrocephalus (iNPH) patients, involving CSF disturbance (28). Although the pathological role of fibrinogen in AD and CAA is increasingly recognized, the mechanisms underlying its passage across the blood-brain barrier (BBB) and its contribution to cerebrovascular damage are still unclear. One potential mechanism is caveolae-mediated transcytosis, a process through which plasma proteins cross endothelial cells (ECs). Yang *et al.* found an age-dependent increase in non-specific caveolar transcytosis of plasma proteins, including fibrinogen, across the BBB (29). Caveolin-1 (Cav-1), a crucial structural protein component of caveolae, plays a significant role in regulating the transfer of molecules from blood to the basolateral side by influencing caveolae formation (30). Furthermore, hyperfibrinogenemia has been linked to increased Cav-1 expression, and elevated levels of fibrinogen have been shown to enhance caveolae formation (31, 32).

Given the lack of a clear mechanism behind fibrinogen extravasation and its role in exacerbating CAA pathology, we hypothesize that fibrin(ogen) extravasation is facilitated by an age-dependent increase in Cav-1 expression, resulting in the co-deposition of fibrin(ogen) and Aβ in the PVS. This, in turn, may trigger neuroinflammation and disrupt the localization of AQP4 in astrocyte end-feet. The loss of AQP4 polarization could impair glymphatic clearance of Aβ, further exacerbating CAA pathology and contributing to cognitive decline. This study aims to elucidate the mechanisms underlying these processes, providing new insights into the interplay between fibrinogen, AQP4, and cerebrovascular dysfunction in CAA and AD pathology.

## Results

### 1) Increase of fibrin(ogen) extravasation and fibrin(ogen)-Aβ co-deposition in TgSwDI mice

To investigate the relationship between CAA and fibrin(ogen) deposition *in vivo*, we analyzed fibrin(ogen) extravasation and co-deposition of fibrin(ogen) with Aβ in TgSwDI mice, a transgenic mouse model of CAA, compared to WT littermates. These mice express human APP harboring the Swedish (K670N/M671L), Dutch (E693Q), and Iowa (D694N) mutations, resulting in the production of Aβ peptides that include two mutations (Dutch and Iowa) associated with HCAA (33). Despite expressing HCAA-type mutant Aβ, TgSwDI mice predominantly exhibit capillary CAA (Type 1), which closely resembles the pathology observed in sporadic CAA—the more common form seen in late onset AD patients. To gain insight into how these vascular pathologies evolve over time, we assessed fibrin(ogen) and Aβ deposition at both early and late stages of disease progression in young and aged mice.

We analyzed CAA, fibrin(ogen) extravasation, and fibrin(ogen)-Aβ co-deposition in TgSwDI mice and WT littermates at 7, 12, and 18 months of age using immunohistochemistry. Fibrin(ogen) was considered extravasated when more than 60% of its total signal was detected outside blood vessels compared to its lumen. We observed significantly increased level of CAA in thalamus of 7-months-old TgSwDI compared to WT littermates, with no change in cortex and hippocampus (Supplementary Fig. S1A and B). However, at 12 and 18 months, CAA was significantly elevated in both the hippocampus and thalamus of TgSwDI mice compared to WT littermates (Fig. 1A–C).

**Figure 1.**
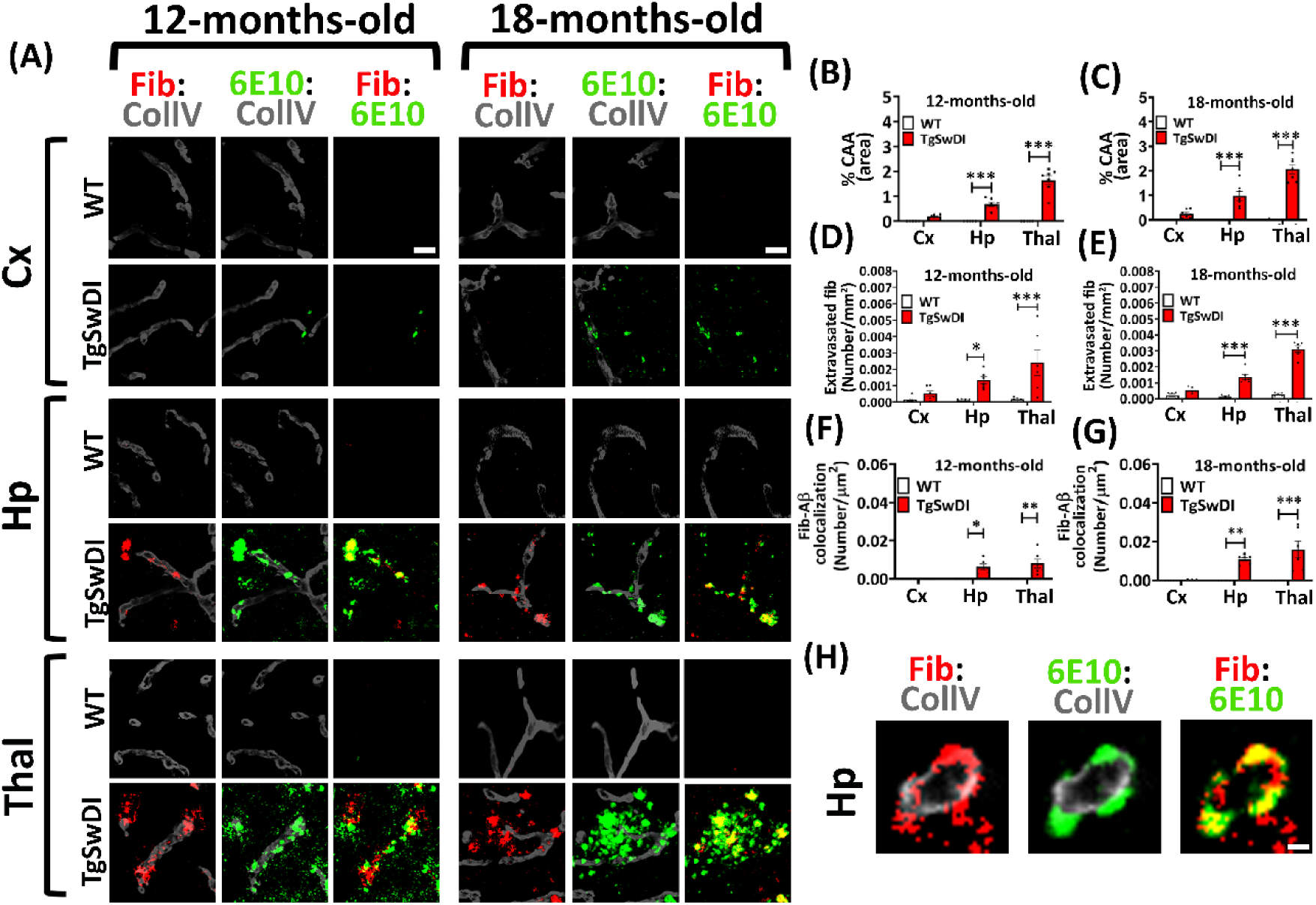
Age-dependent increase of CAA, fibrin(ogen) extravasation and Aβ-fibrin(ogen) co-deposition in the brain of TgSwDI mice. (A) Representative images of brain coronal sections were probed with antibodies against fibrin(ogen) (Fib, red), Aβ (6E10, green), and blood vessels (collagen IV (ColIV), grey). CAA is indicated as colocalized collagen IV and Aβ. Increase of CAA, extravasation of fibrin(ogen), and fibrin(ogen)-Aβ colocalization were observed in hippocampus and thalamus of 12-and 18-months-old TgSwDI compared to WT littermates. Scale=20µm. (B – E) Statistical quantification of % CAA in 12-month-old (B) and 18-months-old (C), number of extravasated fibrin(ogen) spots in 12-months (D) and 18-months-old (E) TgSwDI mice and WT littermates. (F & G) Statistical analysis of fibrin(ogen)-Aβ colocalization in TgSwDI mice compared to WT littermates at both 12-months (F) and 18-months of age (G). (12-months-old mice, n=6 per group; 18-months-old mice, n=5-6 per group). (H) Images of single blood vessel view showed Aβ-fibrin(ogen) co-deposition in perivascular region of hippocampus in 12-months-old TgSwDI mice. Scale=5µm. Data were analyzed by using two-way ANOVA with the Bonferroni post hoc test and shown as average ± SEM; *, P < 0.05, **, P < 0.01, ***P < 0.001. Fib=Fibrin(ogen), ColIV=Collagen IV, Cx=Cortex, Hp=Hippocampus, Thal=Thalamus.

At 7 months, there was no significant extravascular fibrin(ogen) deposition in any brain region of TgSwDI mice compared to WT littermates (Supplementary Fig. S1A and C). In contrast, at 12 and 18 months, we observed marked fibrin(ogen) extravasation in the hippocampus and thalamus of TgSwDI mice (Fig. 1A, D, E). Similarly, significant increase in fibrin(ogen)–Aβ co-deposition were detected at both ages compared to WT controls (Fig. 1A, F, G). High-resolution images of individual vessels in 12-month-old TgSwDI mice revealed clear evidence of fibrin(ogen) leakage and its perivascular co-localization with Aβ (Fig. 1H). It was clearly observed that 18-months-old TgSwDI showed higher CAA, fibrin(ogen) extravasation and fibrin(ogen)-Aβ colocalization than 12-months and 7-months-old mice (Fig.1A-G). Since the hippocampus and thalamus were the brain areas most affected by CAA and fibrin(ogen) extravasation, while the cortex was less impacted in TgSwDI mice, we focused our subsequent analyses on these two regions.

### 2) Perivascular AQP4 loss is associated with fibrin(ogen)-Aβ co-deposits in CAA pathology

Since fibrin(ogen)-Aβ co-deposits are localized to PVS in both HCAA patients (27) and TgSwDI mice, and increased levels of extravasated fibrin(ogen) are associated with reduced perivascular AQP4 in iNPH patients (28), we aimed to investigate whether fibrin(ogen)-Aβ co-deposition affects AQP4 localization around blood vessels. To achieve this goal, we conducted a comprehensive analysis of perivascular AQP4 levels in TgSwDI mice across different ages compared to WT littermates, i) at 7-months (CAA confined to thalamus), ii) 12-months (CAA in both the hippocampus and thalamus), and iii) 18 months (more severe CAA in the hippocampus and thalamus). Additionally, we compared perivascular AQP4 levels between CAA-laden vessels and non-CAA vessels (those without visible CAA pathology) in TgSwDI mice.

At 7-months of age, while there was a significant increase of CAA in the thalamus of TgSwDI mice (Supplementary Fig. S1A and B), there was no significant difference in perivascular AQP4 levels in the thalamus and other brain regions between TgSwDI and WT littermates (Supplementary Fig. S2A and B). By 12- and 18-months of age, TgSwDI mice exhibited a significant decrease of perivascular AQP4 levels in both hippocampus and thalamus, but not in cortex, compared to WT littermates. This decrease was more pronounced at 18-months than at 12-months in TgSwDI mice (Fig. 2A-C). Additionally, perivascular AQP4 was significantly reduced in CAA-laden vessels compared to non-CAA vessels, in hippocampus and thalamus of 12- and 18-months-old TgSwDI mice. Again, the decrease was more severe in 18-months than 12-months TgSwDI (Fig. 2D and E). Interestingly, the loss of perivascular AQP4 in TgSwDI mice was accompanied by increased level of AQP4 outside blood vessels (non-vascular AQP4) (Fig. 2A), suggesting AQP4 depolarization. Taken together, these findings indicate that perivascular AQP4 levels decreases significantly in the regions affected by CAA and AQP4 depolarization appears to occur after the onset of CAA, with greater reductions observed in CAA-laden vessels. Since AQP4 depolarization occurs in the same brain regions and ages as fibrin(ogen)-Aβ co-deposits (Fig. 1), these outcomes suggest a potential link between perivascular fibrin(ogen)-Aβ co-deposits and AQP4 depolarization, which may aggravate CAA pathology.

**Figure 2.**
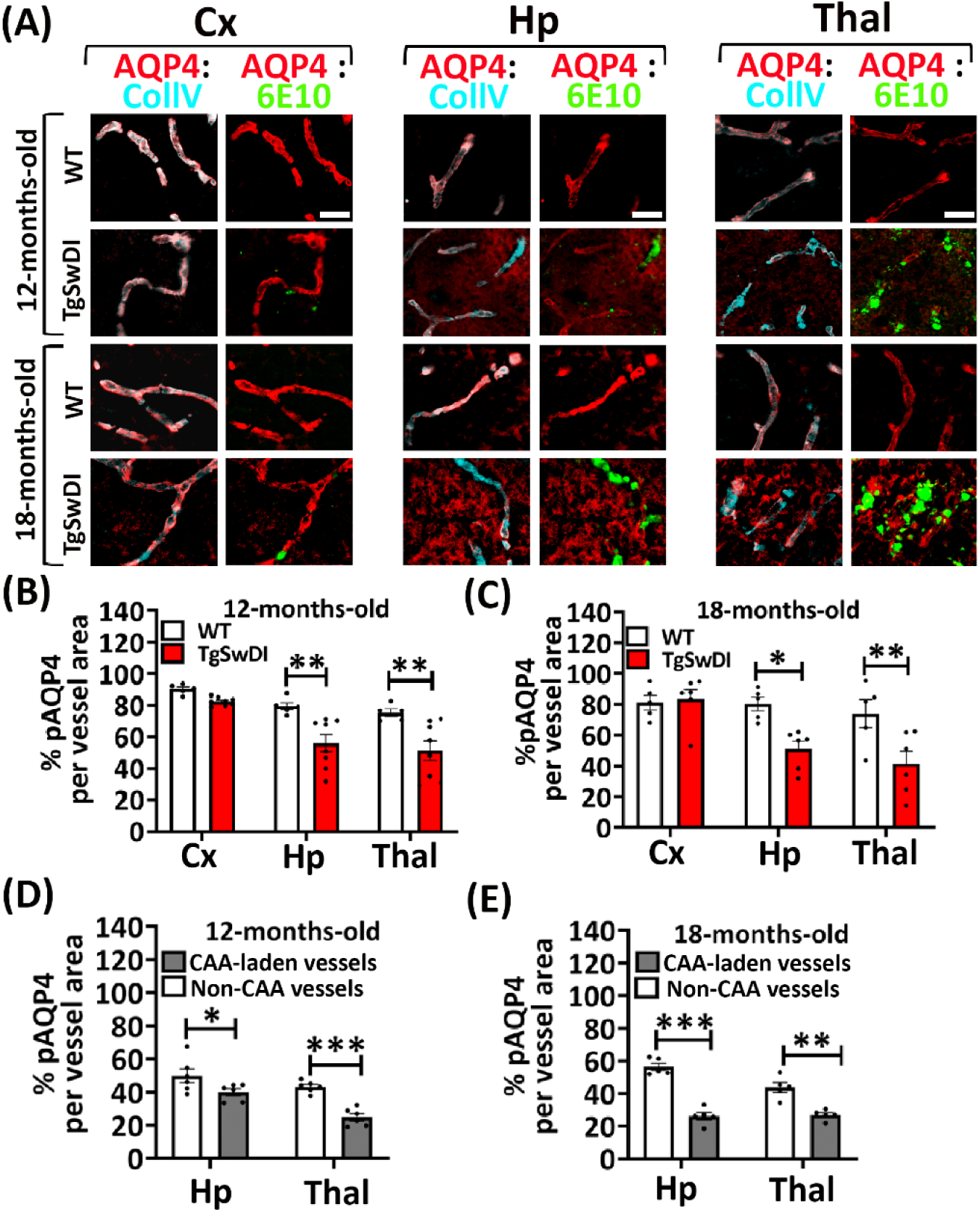
Age dependent decrease of perivascular AQP4 in the brain of TgSwDI mice. (A) Representative images of brain coronal sections of 12- and 18-months-old TgSwDI mice and WT littermates, probed with antibodies against AQP4 (red), Aβ (6E10, green), and blood vessels (collagen IV, cyan). Perivascular AQP4 visualized as pink. Scale=20µm. (B – C) Statistical quantification of perivascular AQP4 in 12- and 18-months-old TgSwDI mice compared to WT littermates. (12-months-old mice, n=5-8 per group; 18-months-old mice, n=5 per group). A significant loss of perivascular AQP4 in the hippocampus and thalamus of 12- and 18-months-old TgSwDI mice was observed. (D -E) Statistical quantification of perivascular AQP4 in CAA-laden vessels compared to non-CAA vessels in 12- and 18-months-old TgSwDI. Perivascular AQP4 in CAA-laden vessels decreased significantly in hippocampus and thalamus, compared to non-CAA vessels in 12- and 18-months-old TgSwDI mice. Data were analyzed by using two-way ANOVA with the Bonferroni post hoc test and shown as average ± SEM; *, P < 0.05, **, P < 0.01, ***P < 0.001. pAQP4=Perivascular AQP4.

### 3) Blood brain barrier integrity is preserved in TgSwDI mice

To explore potential mechanism behind fibrinogen extravasation, we hypothesized that microhemorrhages and BBB disruption could contribute to this process in TgSwDI mice. To test this, we assessed microhemorrhage and BBB integrity in 18-month-old TgSwDI mice and their WT littermates. We first examined microhemorrhages using Prussian blue histology, which detects ferric iron deposits commonly associated with cerebral microbleeds (34). However, we found no significant difference in the number of microhemorrhages in the thalamus of TgSwDI mice compared to WT littermate controls at 18 months of age (Fig. 3A, B). This finding is consistent with observations from other groups (35, 36). To further assess BBB permeability, we administered Evans Blue dye (EBD), a tracer that tightly binds to serum albumin, to 18-month-old TgSwDI and WT mice. The presence of EBD in the brain parenchyma serves as an indicator of BBB leakage (37, 38). Spectrophotometric analysis of brain homogenates from 18-months-old TgSwDI and WT littermates showed no significant differences in EBD content in the cortex and hippocampus (Fig. 3C). To further assess BBB integrity, we analyzed the expression of key tight junction proteins—claudin-5, occludin, and ZO-1—by western blot. The levels of these proteins in the thalamus were comparable between TgSwDI and WT littermates at 18 months of age (Fig. 3D–G). Overall, our findings indicate that microhemorrhages, BBB permeability, and tight junction protein expression are not significantly altered in aged TgSwDI mice, suggesting that these factors are unlikely to explain the observed fibrin(ogen) extravasation in this mouse model of CAA.

**Figure 3.**
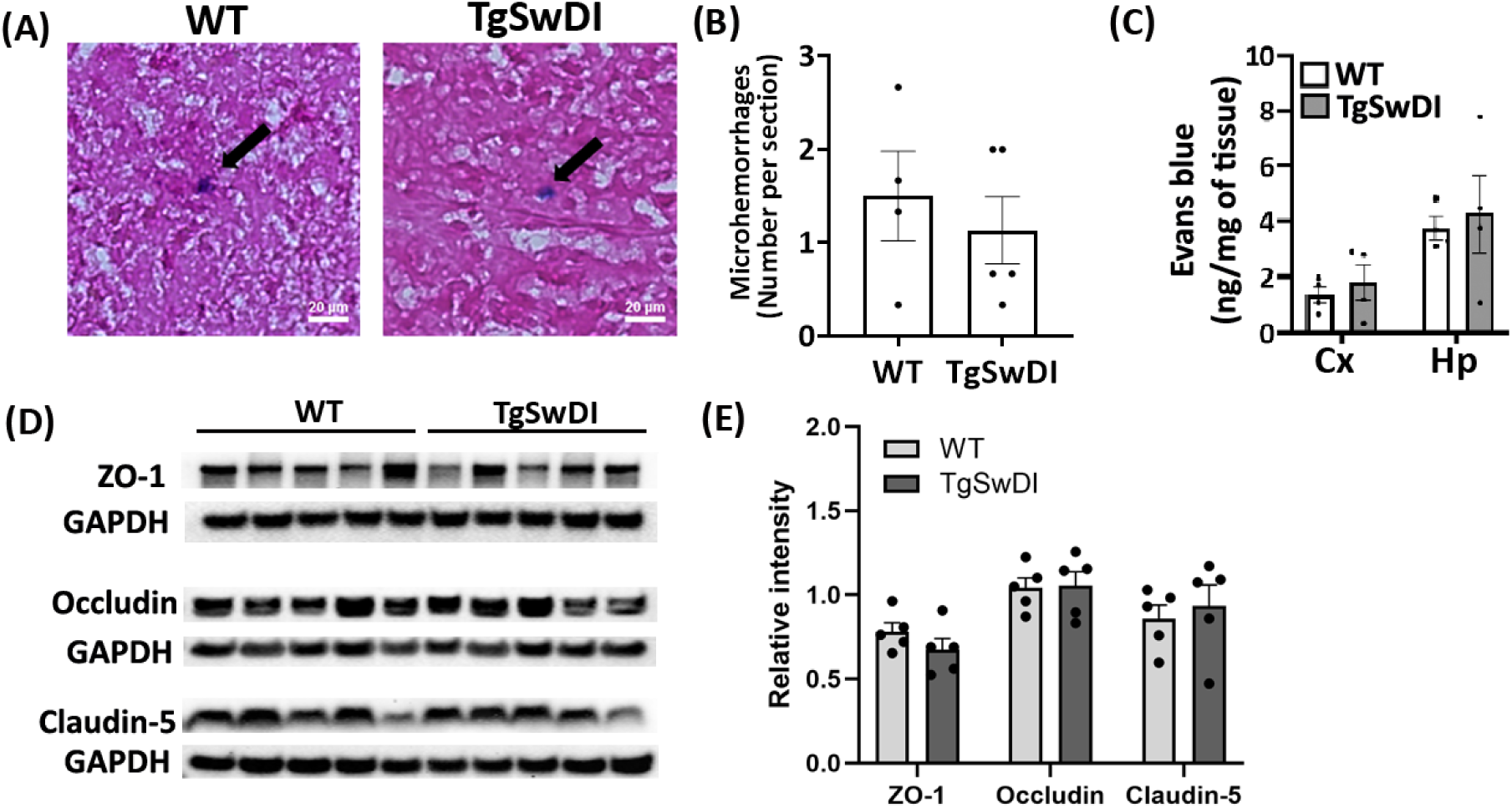
Absence of microhemorrhages and preserved BBB integrity in 18-months-old TgSwDI. (A) There was no significant difference in number of microhemorrhages spots in thalamus between 18-month-old TgSwDI mice and WT littermates. Microhemorrhages were identified by Prussian blue staining (blue color, black arrow). Scale=20µm. (B) Statistical quantification of the number of microhemorrhages spots per tissue section in TgSwDI and WT littermates as observed in (A) (n=4-5 mice per group). (C) Evans blue dye (EBD) levels in lysates from the cortex and hippocampus of 18-months-old TgSwDI and WT mice were determined spectrophotometrically. There was no significant difference of EBD content between TgSwDI and WT littermates (n= 4-5 mice per group). (D) Western blot analysis of tight junction proteins, including ZO-1, Occludin, and Claudin-5, indicated no significant changes in levels of tight junction proteins in the thalamus of 18-months-old TgSwDI mice compared to WT littermates. GAPDH was used as a loading control, with ZO-1 and Occludin sharing the same GAPDH, as they were run on the same gel (n=4-5 mice per group). (E, F, G) Densitometric quantification of the tight junction proteins, as shown in (D). Data were analyzed using Student’s t-test and are presented as mean ± SEM.

### 4) Caveolae-mediated transcytosis could be altered in an age-dependent manner in TgSwDI Mice

Given the preserved BBB integrity in TgSwDI mice, we sought to identify alternative molecular mechanisms that could facilitate the extravasation of fibrinogen across the BBB, leading to its deposition in the PVS. Recent study has found that an age-dependent increase in non-specific caveolae-mediated transcytosis facilitates BBB transport of circulatory plasma proteins including fibrinogen (29). Cav-1 plays a crucial role in compromising BBB permeability, allowing plasma proteins, to cross the BBB via caveolae-mediated transcytosis (39, 40). We hypothesized that Cav-1 levels could be increased in the brains of aged TgSwDI mice, facilitating the transcytosis of fibrinogen across brain endothelial cells and resulting in its deposition in the PVS. To test this, we analyzed Cav-1 levels in endothelial cells by assessing their colocalization with CD31 (an endothelial cell marker) in TgSwDI mice and WT littermates in an age-dependent manner. We found that at 7-months of age, there was no significant difference in the endothelial Cav-1 expression between TgSwDI and WT littermates across brain regions (Supplementary Fig. S3A and B). However, at both 12 and 18-months of age, TgSwDI mice showed significantly higher endothelial Cav-1 level in the hippocampus and thalamus, compared to WT littermates (Fig. 4A-C). To ensure unbiased analysis and confirm the age dependent Cav-1 expression changes, we analyzed whole hemisphere of coronal sections of TgSwDI mice and WT littermates at ages 7-, 12- and 18-months. WT mice showed significant age-dependent increase of Cav-1 in the cortex and TgSwDI mice exhibited age-dependent increase of Cav-1 in all three regions (Supplementary Fig. S4A-D). The age-dependent increase in endothelial Cav-1 levels in the hippocampus and thalamus of aged TgSwDI mice suggests that these molecular changes may enhance caveolae-mediated transcytosis, contributing to fibrinogen extravasation across the BBB and implicating Cav-1 upregulation as a key driver of BBB permeability, potentially exacerbating vascular pathology in TgSwDI mice.

**Figure 4.**
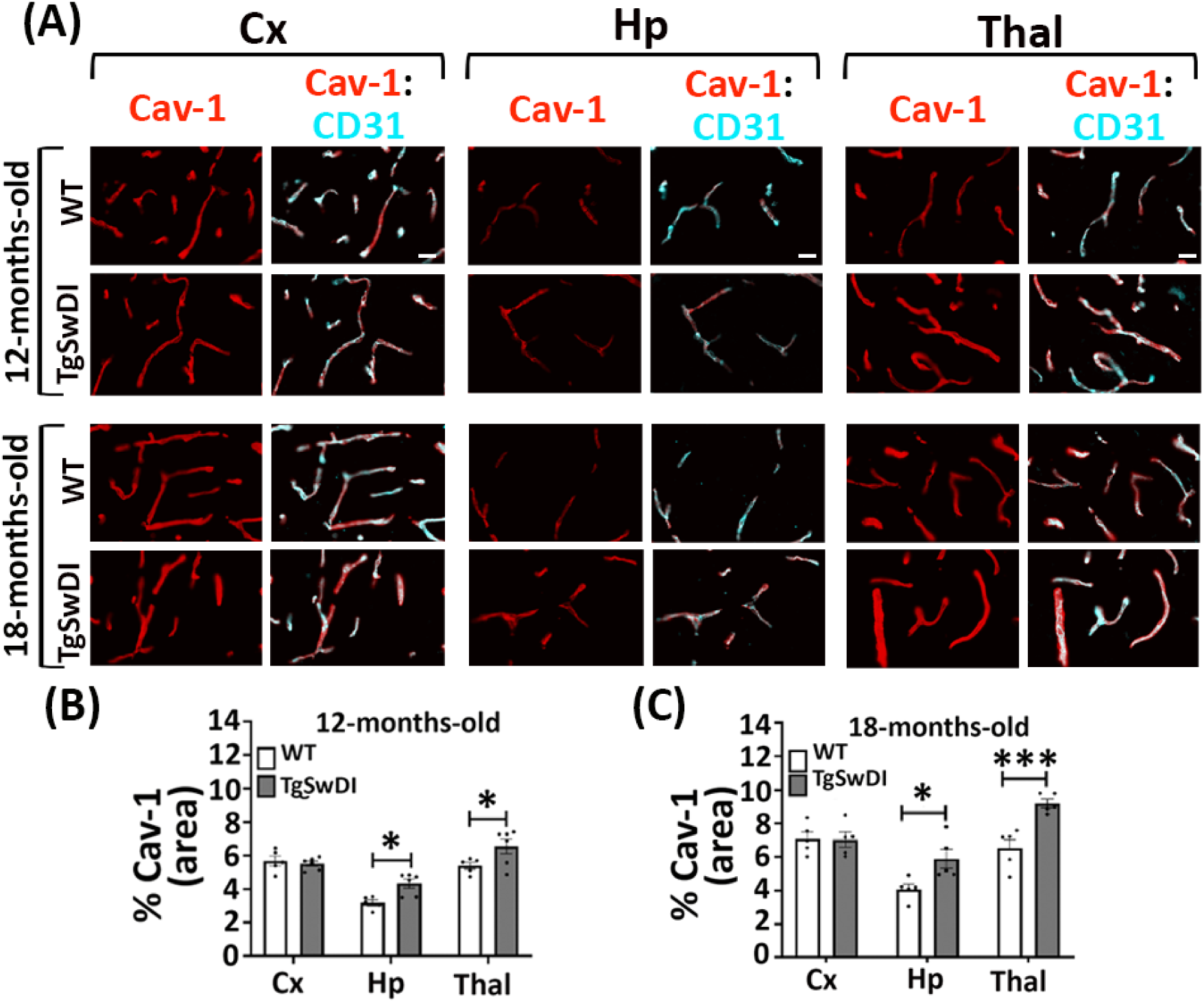
Altered Cav-1 expression in the brain of TgSwDI mice. (A) Representative images of coronal sections of TgSwDI mice and WT littermates at 12-months and 18-months-old were probed with antibodies against Cav-1 (red) and endothelial cell marker-CD31 (cyan). (B, C) Statistical quantification of (A) showed significant increase of Cav-1 in hippocampus and thalamus of 12- and 18-months-old TgSwDI mice compared to WT littermates (12-months-old mice, n=5-6 per group; 18-months-old mice, n=5 per group, three tissue sections per mice). Scale=20µm. Data were analyzed using two-way ANOVA with the Bonferroni post hoc test and shown as average ± SEM; *, P < 0.05, **, ***P < 0.001.

### 5) Depletion of plasma fibrinogen decreases CAA and restores impaired spatial memory in TgSwDI mice

Given the potential link between fibrin(ogen) extravasation, CAA and other CAA-related pathologies, we explored the impact of fibrinogen depletion on these pathological features in TgSwDI mice. Ancrod, a defibrinogenating agent derived from the venom of the Malayan pit viper, has previously been reported to reduce fibrinogen-mediated pathology by lowering fibrinogen levels in the blood, thereby decreasing CAA pathology and improving memory in AD mice (23). However, clinical study for ischemic stroke showed that ancrod increased the risk of intracerebral hemorrhage, and its efficacy has not been clearly demonstrated, highlighting the need for alternative approaches (41). To address this, we utilized lipid nanoparticle (LNP)-encapsulated small interfering RNA (siRNA) specifically targeting the fibrinogen alpha chain (siFga) to deplete fibrinogen in the blood. Since both bleeding time and blood loss were comparable between siFga-treated and control mice, along with the absence of bleeding risk in siFga-treated mice (42), this siRNA approach is unlikely to significantly impact hemostasis. This approach has been used safely in various animal models, leading to robust and controllable depletion of plasma fibrinogen (43–45). In our study, we administered siRNA targeting luciferase (siLuc) as a negative control. For clarity and consistency throughout the manuscript, mice treated with siLuc will be referred as siLuc WT or siLuc TgSwDI, and those treated with siFga will be designated as siFga WT or siFga TgSwDI.

Mice were sacrificed at 12-months of age following three months of treatment with siRNA LNPs. Western blot analysis confirmed the efficacy of siFga, showing a reduction in plasma fibrinogen levels by more than 90% compared to siLuc-treated mice (Fig. 5A and B). While siLuc TgSwDI mice exhibited significantly higher CAA in the hippocampus and thalamus compared to siLuc WT and siFga WT mice, a significant reduction in CAA was observed in the hippocampus and thalamus of siFga-treated TgSwDI mice compared to siLuc TgSwDI mice (Fig. 5C and D). We further analyzed whether fibrinogen depletion affected Aβ plaque pathology and found that siFga treatment did not alter Aβ plaque levels in any of these brain regions in siFga TgSwDI mice (Supplementary Fig. S5A and B). These results suggest that fibrinogen depletion specifically reduces CAA pathology around blood vessels without affecting Aβ plaques in the brain parenchyma.

**Figure 5.**
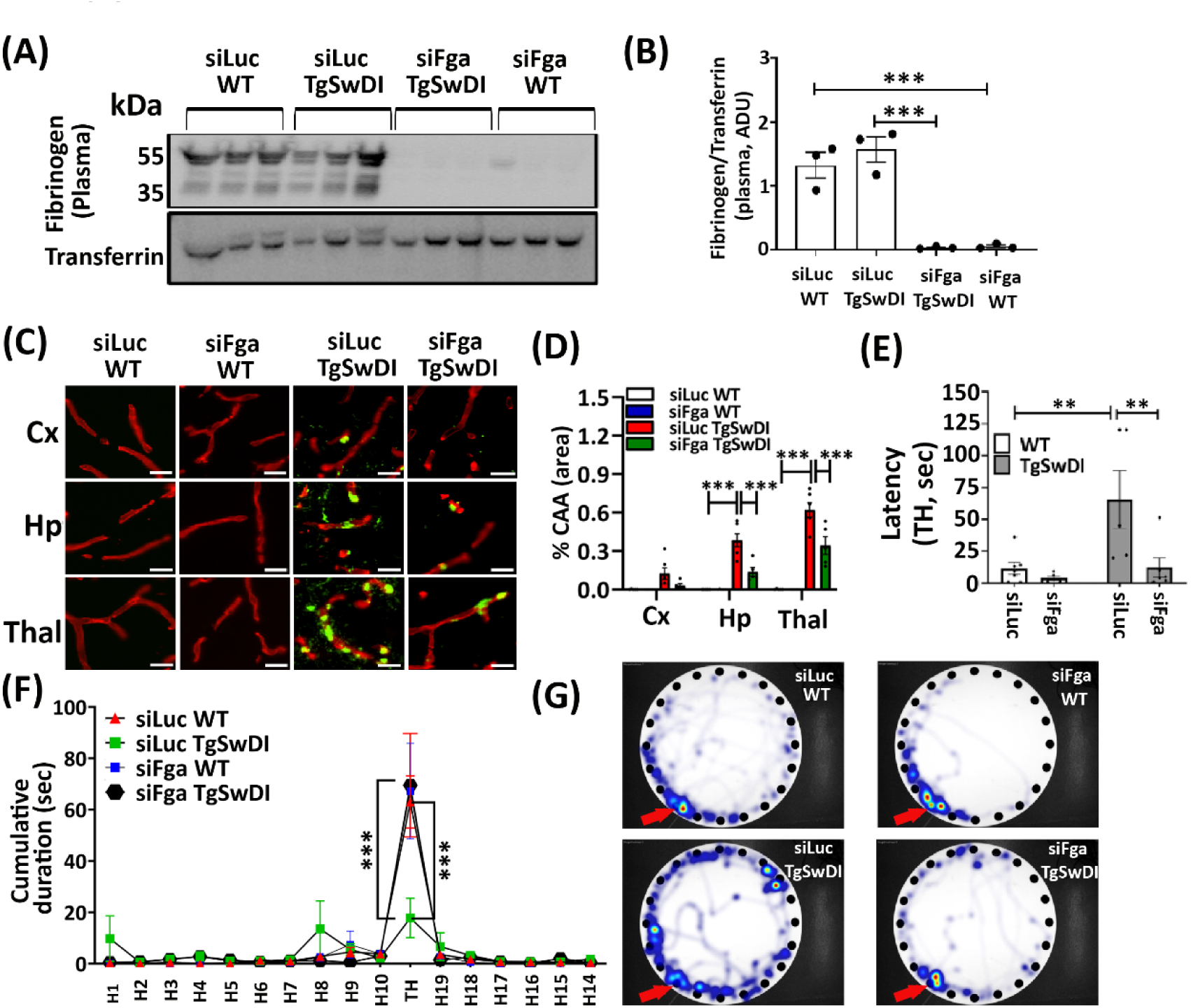
Depletion of fibrinogen significantly reduced CAA and restored spatial memory in TgSwDI mice. (A) Western blot analysis confirmed fibrinogen depletion in plasma isolated from siRNA-LNPs treated mice. Transferrin was used as a loading control (n=3 per group). (B) Densitometric quantification of the western blot from (A), illustrating a significant reduction of plasma fibrinogen levels. (C) Brain sections were immunostained with 6E10 to detect Aβ (green) and anti-collagen IV to identify blood vessels (red). Scale=20µm. (D) Statistical quantification of (C) showing significant decrease of CAA in the hippocampus and thalamus of siFga TgSwDI mice compared to siLuc-treated TgSwDI mice. (n= 5-6 mice per group, three tissue sections per mice). (E & F) Spatial memory of siRNA-treated mice was assessed using the Barnes maze task (n= 5-7 mice per group). During the probe trial, siFga TgSwDI mice showed a decreased latency to reach the target hole (E) and spent significantly more time (cumulated duration) around the target hole (TH) compared to siLuc TgSwDI mice (F). Hole numbers such as H1 and H2, denote the respective hole locations, with TH indicating the target hole. (G) The heatmap illustrates the movement paths and the duration spent in different areas of the Barnes maze, using a color gradient. Regions where the mice spent more time are represented in warm colors (red/yellow), while regions with less time spent are shown in cooler colors (blue). The heatmap displays the average duration spent in area for each group of mice, with the red arrow pointing to the target hole. Data were analyzed by using two-way ANOVA with the Bonferroni post hoc test and shown as average ± SEM; **, P < 0.01, ***P < 0.001.

Given the association between CAA and cognitive decline, and the reported spatial memory deficits in TgSwDI mice as assessed by the Barnes maze task (46–48), we investigated whether fibrinogen depletion could improve spatial memory impairment in these mice. Upon fibrinogen depletion, siFga-treated TgSwDI mice exhibited significant improvements in spatial memory compared to siLuc-treated TgSwDI mice, as demonstrated by the Barnes maze task (Fig. 5E-G). During the probe trial, siFga-treated TgSwDI mice showed a significantly reduced latency to locate the target hole (TH) compared to siLuc-treated TgSwDI mice, reaching levels similar to siLuc WT mice (Fig. 5E). Additionally, the cumulative time spent in the TH was significantly higher in siFga TgSwDI mice than in siLuc TgSwDI mice, restoring it to levels comparable to siLuc WT mice (Fig. 5F).

To further visualize spatial search strategies, we generated heatmaps, displaying movement paths and the duration spent in different areas of the Barnes maze as a color gradient. Areas where mice spent more time appear as warm colors (red/yellow), while those with shorter durations appear as cool colors (blue). The heatmaps revealed that siLuc TgSwDI mice spent less time near the TH compared to siLuc WT and siFga TgSwDI mice, indicating impaired spatial memory (Fig. 5G) (Supplementary Fig. S6A). Notably, siFga WT mice exhibited spatial memory performance comparable to siLuc WT mice across all analyzed parameters, suggesting that fibrinogen depletion does not induce cognitive deficits in WT mice. We also evaluated locomotor activity and anxiety-like behavior in the animals using the open field test. Analysis of total distance traveled, and the time spent in the center (a measure of anxiety) revealed no significant differences between any of the siRNA-treated groups (Supplementary Fig. S6B and C). Overall, these findings suggest that fibrinogen depletion alleviates CAA-associated spatial memory deficits in TgSwDI mice without impacting locomotor function or anxiety-like behavior.

### 6) Fibrinogen depletion restored perivascular AQP4 localization and decreased Cav-1 levels in TgSwDI mice

Given the possibility that perivascular fibrin(ogen)-Aβ co-deposits may be associated with AQP4 depolarization (Fig. 2), we examined whether fibrinogen depletion, linked to a reduction in CAA, could restore perivascular AQP4 polarization in TgSwDI mice. Our results showed that siFga-treated TgSwDI mice exhibited significantly higher perivascular AQP4 levels in both the hippocampus and thalamus compared to siLuc TgSwDI mice, with levels similar to siLuc WT (Fig. 6A and B). In contrast, siLuc-treated TgSwDI mice exhibited significantly lower perivascular AQP4 levels compared to siLuc WT (Fig. 6A and B). Simultaneously, non-vascular AQP4 levels were significantly reduced in siFga TgSwDI mice compared to siLuc TgSwDI mice in both hippocampus and thalamus (Fig. 6A and C) (Supplementary Fig. S7). These findings suggest that fibrinogen depletion can restore perivascular AQP4 levels in TgSwDI mice, which may further contribute to reducing CAA by enhancing Aβ clearance.

**Figure 6.**
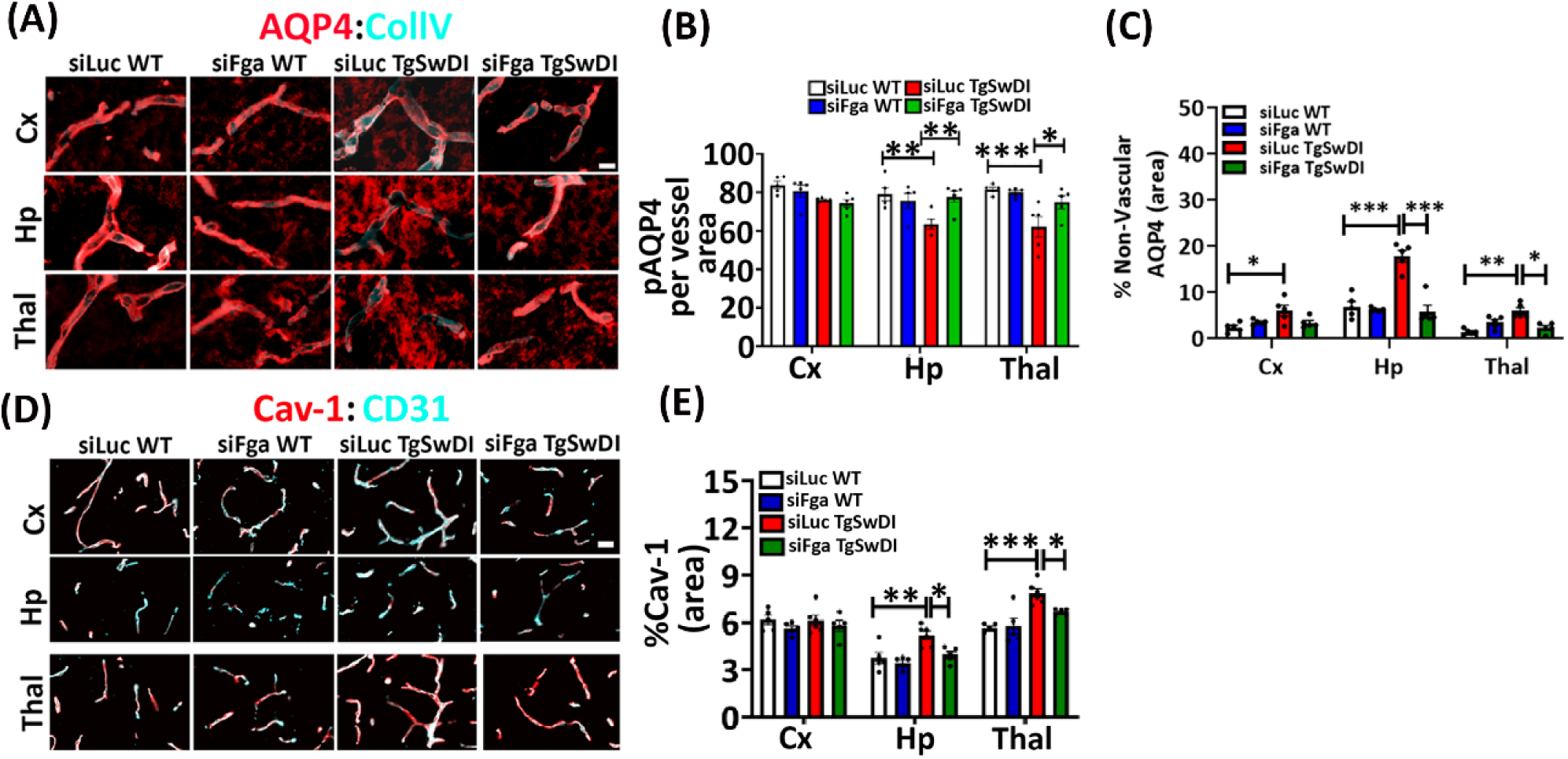
Fibrinogen depletion restores perivascular AQP4 and Cav-1 levels in TgSwDI mice to levels similar to WT littermates. (A) Brain sections were immunostained for AQP4 (red) and Collagen IV (cyan). An increase in perivascular AQP4 was found in hippocampus and thalamus of siFga TgSwDI compared to siLuc TgSwDI. siLuc TgSwDI showed lower perivascular AQP4 levels as compared to siLuc WT in both hippocampus and thalamus. Scale=10µm. (B) Statistical quantification of perivascular AQP4 levels as observed in (A) (n=5 mice per group). (C) Statistical quantification of non-vascular AQP4 levels as observed in (A) (n=5 per group, three tissue sections per mice). (D) Coronal brain sections were immunostained with Cav-1 (red) and CD31(cyan). A decrease of Cav-1 was observed in both hippocampus and thalamus of siFga TgSwDI as compared to siLuc TgSwDI. siLuc TgSwDI mice exhibited significantly higher Cav-1 levels compared to siLuc WT mice in both the hippocampus and thalamus. Scale=20 µm. (E) Statistical quantification of Cav-1 levels as observed in (D) (n=5-6 mice per group). Data were analyzed by using two-way ANOVA with the Bonferroni post hoc test and shown as average ± SEM; *, P < 0.05, **, P < 0.01, ***P < 0.001.

A previous study reported that in a mouse model of hyperfibrinogenemia, a condition characterized by elevated plasma fibrinogen levels, both Cav-1 expression and caveolae-mediated transcytosis are significantly increased (31). Based on this, we investigated the impact of fibrinogen depletion on Cav-1 expression in TgSwDI mice. We found that siFga-treated TgSwDI mice exhibited significantly lower Cav-1 expression in the hippocampus and thalamus compared to siLuc TgSwDI mice. As expected, siLuc TgSwDI showed significantly higher Cav-1 expression compared to siLuc WT in both the regions (Fig. 6D and E). These results suggest that reducing blood fibrinogen levels could decrease endothelial Cav-1 expression in TgSwDI mice.

### 7) Fibrinogen depletion mitigates microglial-mediated neuroinflammation in CAA affected regions of TgSwDI mice

Fibrin(ogen) has been associated with microglial activation via the Mac-1 integrin receptor, where CD11b serves as the α chain of Mac-1 (49). Fibrin(ogen) triggers rapid microglial responses toward the vasculature and plays a key role in axonal damage during neuroinflammation (50). Given fibrin(ogen) is proinflammatory, and its colocalization with Aβ in the PVS contributes to persistent neuroinflammation, we investigated the impact of fibrinogen depletion on microglia We used CD11b as a marker for microglial cells and observed a significant increase in ameboid microglial morphology, indicating activation, in the cortex, hippocampus, and thalamus of siLuc TgSwDI mice compared to siLuc WT (Fig. 7A). In siFga TgSwDI mice, activated microglia were significantly reduced in the hippocampus and thalamus but remained unchanged in the cortex (Fig. 7B). Notably, in siLuc TgSwDI mice, activated microglia in the hippocampus and thalamus were primarily located around CAA and blood vessels, whereas in the cortex, they were mainly associated with Aβ plaques (Fig. 7A). Whole-brain coronal sections further confirmed these regional differences in CD11b expression (Supplementary Fig. S8).

**Figure 7.**
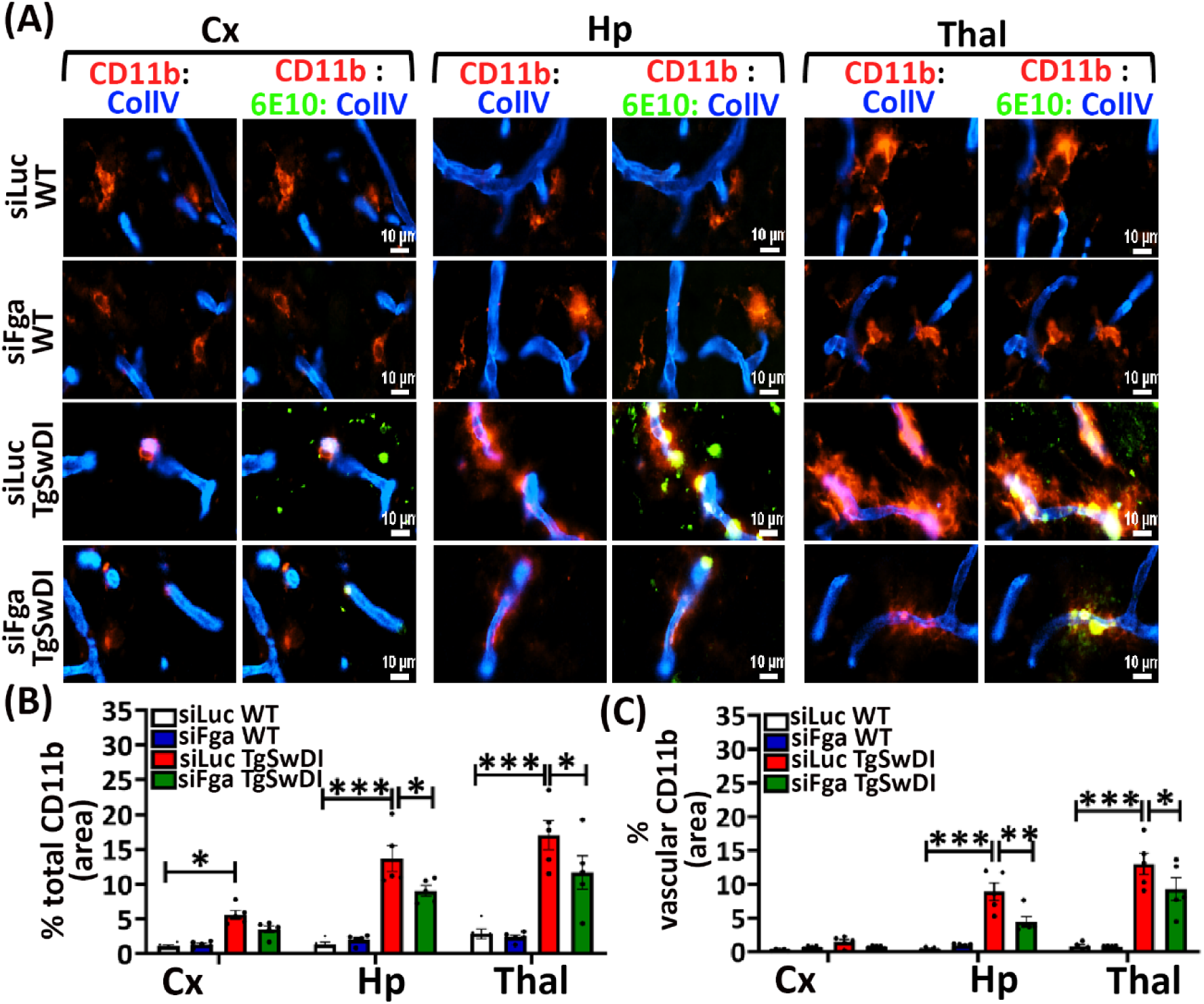
Fibrinogen depletion reduces microglial activation associated with CAA laden vessels in TgSwDI mice. (A) Coronal brain sections immunostained for CD11b (red) representing microglia, Aβ (6E10, green) and collagen IV (blue) showed decrease in activated microglial cells in the hippocampus and thalamus of siFga TgSwDI mice compared to siLuc TgSwDI mice. siLuc TgSwDI mice exhibited significantly higher activated microglial cells compared to siLuc WT mice in cortex, hippocampus and thalamus. Scale=10µm. Statistical quantification of the area of microglial cells (B) and microglial cells near blood vessels (vascular CD11b) (C) was performed across the cortex, hippocampus, and thalamus as observed in (A) (n=5 mice per group). Data are expressed as mean ± SEM and analyzed using two-way ANOVA with Bonferroni post hoc test; *P < 0.05, **P < 0.01, ***P < 0.001.

The selective reduction of activated microglia in the hippocampus and thalamus likely reflects the targeted effect of fibrinogen depletion on CAA pathology, which is more prominent in these regions. Since siFga treatment effectively reduced CAA in these areas without altering Aβ plaque levels, the observed decrease in microglial activation is likely linked to CAA reduction rather than Aβ pathology. In contrast, cortical microglial activation remained unchanged, as fibrinogen depletion did not significantly affect Aβ plaques, the primary trigger for microglial activation in this region. To further investigate vascular microglia, we analyzed their activation around blood vessels and found a significant increase in siLuc TgSwDI mice compared to siLuc WT in the hippocampus and thalamus, but not in the cortex (Fig. 7C). siFga treatment significantly reduced vascular microglial activation in these regions (Fig. 7C). These results suggest that fibrinogen depletion specifically reduces vascular microglial activation around CAA, where fibrinogen and Aβ co-deposit.

TgSwDI mice are also known to exhibit pronounced GFAP-positive reactive astrocytes (51). Therefore, we further analyzed whether fibrinogen depletion affects GFAP-positive reactive astrocytes. We found a significant increase in GFAP expression in the cortex and thalamus of siLuc TgSwDI mice compared to siLuc WT mice. However, there were no significant differences in GFAP immunoreactivity between fibrinogen-depleted TgSwDI mice and those with normal blood fibrinogen levels, suggesting that fibrinogen depletion does not significantly impact reactive astrocyte populations in TgSwDI mice (Supplementary Fig. S9).

## Discussion

In this study, we present a potential mechanism by which fibrin(ogen) extravasation may exacerbate cerebrovascular pathology, including CAA, neuroinflammation, and AQP4 depolarization. Our results using siRNA-LNPs suggest that the CAA-associated cerebrovascular pathology in TgSwDI mice may be driven by two interrelated mechanisms (Supplementary Fig. S10). First, an age-dependent increase in Cav-1-mediated transcytosis facilitates fibrinogen extravasation into the PVS. The extravasated fibrin(ogen) binds to Aβ peptides, promoting fibrin(ogen) deposition, which further amplifies Cav-1 expression in a feedback loop that exacerbates the pathology. Simultaneously, fibrin(ogen)-Aβ co-deposits in the PVS activate microglia and may cause the depolarization of AQP4, further intensifying CAA formation. These findings underscore the complex interplay between Cav-1, fibrin(ogen) extravasation, and AQP4 depolarization, providing deeper insights into the progression of CAA.

In this study, we examined the progression of cerebral amyloid angiopathy (CAA) and related vascular pathologies in TgSwDI mice at three time points—7, 12, and 18 months of age— representing early and advanced stages of disease. At 7 months, TgSwDI mice exhibited early CAA pathology, primarily localized to the thalamus, but showed no evidence of fibrin(ogen) extravasation (Supplementary Fig. S1), AQP4 depolarization (Supplementary Fig. S2), or changes in Cav-1 expression (Supplementary Fig. S3). By 12 and 18 months, the disease had progressed, with marked increases in CAA burden, fibrin(ogen) extravasation, and fibrin(ogen)– Aβ co-deposition in both the hippocampus and thalamus (Fig. 1), accompanied by AQP4 depolarization (Fig. 2) and upregulation of Cav-1 (Fig. 4). These findings suggest that vascular Aβ deposition is the earliest detectable pathological event in TgSwDI mice, emerging by 7 months of age with the thalamus as the initial site of CAA development. CAA occurs prior to fibrin(ogen) extravasation, AQP4 depolarization, or Cav-1 upregulation, indicating that amyloid deposition precedes other vascular dysfunctions in the progression of disease in this model.

Regulation of transcytosis is proposed as a critical mechanism for maintaining selective permeability of BBB in CNS (52, 53). Cav-1 mediated transcytosis plays an important role in compromising BBB integrity (39, 40). Brain tissues from AD patients show increased levels of Cav-1 in the hippocampus and frontal cortex, relative to age-matched control subjects (54). Our results showed significantly higher expression of endothelial Cav-1 in the hippocampus and thalamus of TgSwDI mice in an age dependent manner (Fig.4; Supplementary Fig. S4), suggesting that age may contribute to enhanced Cav-1 expression in these brain areas, as previously reported (55, 56). These brain regions also exhibit significant fibrin(ogen) extravasation at same ages (Fig. 1) indicating a temporal and spatial correlation between Cav-1 overexpression and the leakage of fibrin(ogen). Therefore, our results suggest that enhanced endothelial Cav-1 expression in TgSwDI mice may facilitate the leakage of fibrinogen into the perivascular area through non-specific caveolar transcytosis. Apart from our findings suggesting Cav-1’s potential role in fibrinogen extravasation, there are reports suggesting that fibrinogen could also enhance endothelial Cav-1 levels. In a mice model of hyperfibrinogenemia, Muradashvili et al., demonstrated that elevated blood levels of fibrinogen alter cerebrovascular permeability mainly by increasing caveolae-mediated transcytosis (31) and elevating Cav-1 expression (27). As fibrinogen binding to Aβ forms fibrinolysis resistant clot (19), once fibrinogen extravasates and binds to Aβ, it can persist in the PVS potentially leading to enhanced Cav-1 expression. Thus, our study demonstrating an age-dependent increase in Cav-1 levels in TgSwDI mice, along with a decrease of Cav-1 levels in siFga-treated TgSwDI mice, suggests that the accumulation of fibrin(ogen) in the PVS may also stimulate Cav-1 upregulation in endothelial cells, in addition to the increase driven by aging.

Our study also showed a strong association between fibrin(ogen)-Aβ co-deposition and AQP4 depolarization. This finding is supported by several previous reports indicating the relationship between AQP4 with Aβ clearance, CAA pathology, and fibrinogen extravasation. *Aqp4* gene deletion and depolarization of AQP4, reduce Aβ clearance and promote microvascular Aβ deposition (13, 14, 57–59). Additionally, altered expression and delocalization of AQP4 have been documented in AD and CAA patients across several reports (18–21). Notably, fibrinogen extravasation has been associated with decreased perivascular AQP4 levels in iNPH patients, leading to impaired clearance of brain metabolic waste and increased Aβ plaque burden, similar to what is observed in AD patients (28, 60). In our study, we observed that as CAA spread to the hippocampus and thalamus in aging TgSwDI mice (12 and 18 months), there was a progressive loss of perivascular AQP4 in both regions (Fig. 2). Importantly, CAA-positive vessels showed significantly lower levels of perivascular AQP4 compared to CAA-free vessels (Fig. 2). Furthermore, fibrinogen depletion in siFga-treated TgSwDI mice not only reduced CAA burden (Fig. 5) but also restored AQP4 polarity in both the hippocampus and thalamus (Fig. 6). Together, these findings suggest that fibrin(ogen) accumulation contributes to CAA-associated AQP4 depolarization, and that restoring perivascular AQP4 localization may represent a potential therapeutic strategy to improve Aβ clearance and limit disease progression.

Our study elucidates the mechanisms by which fibrin(ogen)-Aβ colocalization in PVS led to cerebrovascular dysfunctions, highlighting the critical role of fibrin(ogen) in CAA pathology. We propose that the age-dependent increase in Cav-1 expression in TgSwDI mice may contribute to fibrin(ogen) extravasation. This extravasation allows fibrinogen to bind to Aβ, triggering microglial neuroinflammation and AQP4 depolarization, which worsens CAA pathology (Supplementary Fig. S10). Fibrin(ogen) knockdown not only reduced CAA pathology but also improved cognitive function, suggesting the therapeutic potential of targeting fibrin(ogen) in CAA-related pathologies.

## Materials and Methods

### Animals

TgSwDI (The Jackson Laboratory, stock #034843-JAX) are transgenic mice on C57BL/6 background that express human APP gene (isoform 770) containing the Swedish (K670N/M671L), Dutch (E693Q), and Iowa (D694N) mutations under the control of the mouse Thy1 promoter (33). The Swedish K670N/M671L mutation results in enhance β-secretase processing and increased production of Aβ. Dutch and Iowa mutations increase CAA in the brain of TgSwDI mice (33). TgSwDI hemizygous transgenic mice and WT littermate controls were used in the entire study. The mice were housed under a 12-hour light/dark cycle and had ad libitum access to food and water. All work involving the maintenance, breeding, and experimentation with animals adhered to protocols approved by the Rutgers University Institutional Animal Care and Use Committee. All behavioral and imaging analyses were conducted blind to genotype and treatment to minimize potential bias.

### Intravenous administration of siRNA lipid nano particles

A previously identified 2’*O*-methylated siRNA sequence targeting murine fibrinogen (*Fga)* mRNA was used (42) and an siRNA targeting luciferase (siLuc) was applied as a negative control. Sequences were obtained commercially by Integrated DNA Therapeutics (IDT) and siRNAs were encapsulated in lipid nanoparticle (LNP) as previously described (61). Briefly, siRNAs were dissolved in sodium acetate (pH 4) and combined with a lipid solution at an amine-to-phosphate (N/P) ratio of 3. The LNP consisted of an ionizable cationic lipid, DLin-MC3-DMA, and 3 helper lipids: cholesterol, DSPC and PEG-DMG at a 50: 38.5:10:1.5 % molar ratio (Avanti Lipids). The LNP were dialyzed against Dulbecco’s phosphate buffered saline (PBS) at pH 7.4 in 500-fold volume excess. Cholesterol content was measured using the Cholesterol E Assay Kit (Wako Chemicals, Mountain View, CA USA) and RiboGreen assay (Quant-IT Ribogreen RNA Assay Kit, ThermoFisher) was performed to determine mRNA concentration and encapsulation efficiency. siRNA-LNP were diluted to 0.2 mg/mL with a buffer containing 10 mM L-histidine and 10% sucrose (pH 7.4) and stored at −80°C. 9 months-old TgSwDI mice and WT littermates were intravenously injected with LNP-encapsulated siRNA targeting the fibrinogen alpha chain (siFga) and siRNA sequence targeting luciferase (siLuc) as a negative control, every 10 days at a dose of 2 mg of siRNA per kg of body weight for three months to maintain the knockdown.

#### Immunohistochemistry

TgSwDI mice at the age of 7-or 12- or 18 months, or mice administered with siRNA-LNPs were perfused transcardially with 0.9% saline plus 2.5% heparin solution, followed by extraction of brains and embedding them in optical cutting temperature (OCT) medium. Brains were frozen and then sectioned (20μm) coronally, using cryostat (Leica CM1950). Sections spanning bregma -2.0 to -2.5 mm were mounted on glass slides, followed by immunohistochemistry.

Sections were fixed with chilled methanol: acetone (1:1) for 5 minutes or 4 % paraformaldehyde for 20 minutes at room temperature. Sections were then washed with PBS and then blocked in PBST (PBS+0.3% Tritonx-100) with 5% normal donkey serum for 1 hour at room temperature in humidified chamber. Thereafter, sections were incubated with primary antibody overnight diluted in PBST solution (PBS+0.1% TritonX-100) with 5% normal donkey serum at 4°C in humidified chamber. The primary antibody used were 1) anti-amyloid-beta, 1-16 (6E10) conjugated to Alexa Fluor 488 (1/250) (BioLegend, 803013), 2) anti-fibrinogen (1/500) (Dako, A008002-2), 3) anti-collagen IV (1/250) (Millipore, AB769), 4) anti-Caveolin-1 (1/1000) (Cell signalling technology, 3267S), 6) anti-CD31 (1/30) (BD Pharmingen, 550274), 7) anti-CD11b (1/200) (Cell signalling technology, 17800), 8) anti-GFAP (1/250) (Thermofisher scientific, 3-0300), 9) anti-AQP4 (1/500) (Cell signalling technology, 59678S). Sections were then washed 3 times with PBST (PBS+0.05% TritonX-100) for 5 minutes on shaker. Thereafter, sections were incubated with secondary antibodies, diluted in PBST (PBS+0.1% TritonX-100) with 5 % donkey serum for 1 hour at room temperature in humidified chamber. The secondary antibodies were tagged with Alexa Flour 405 or 488 or 555 or 647 fluorophores. The sections were washed 3 times with PBST (PBS+0.05% TritonX-100) for 5 minutes on shaker. The sections were incubated with Sudan black b (Sigma, 199664) for 1 minute to diminish lipofuscin autofluorescence and then washed with 70% ethanol for 20 seconds, followed by washing with PBS, 3 times for 5 minutes on shaker. The sections were then mounted with Vectashield® antifade mounting medium (Vectorlabs, H-1000-10) and cover slipped.

For fibrin(ogen)-Aβ co-deposition imaging, Nikon A1R confocal microscope was used, while other images were acquired using Nikon eclipse Ti2 fluorescence microscope equipped with a 20x/0.75 NA objective with NIS elements AR software. Images were acquired using a 20x/0.75 NA objective at a 16-bit depth. All images were captured by tiling the entire coronal section with the same exposure and gain settings to ensure an unbiased comparison between TgSwDI and WT littermates. Three different sections per mouse were analyzed using Fiji-ImageJ software. Vascular and total CD11b areas were quantified in siRNA-LNP-treated mice using an automated, unbiased image processing approach with the MATLAB Image Processing Toolbox. CD11b signals were taken as vascular, if 10% or more colocalize with Collagen IV signals. To assess this, we created a MATLAB code (Supplementary File 1) which determines boundary boxes around continuous objects in the CD11b staining channel that were larger than 20 square microns. For each continuous object selected, we calculated the area of CD11b staining as well as area of collagen IV staining. Using these two parameters, we were able to determine which areas of CD11b fell into the category of vascular CD11b staining. MATLAB was also utilized for analyzing perivascular AQP4 levels in CAA and non-CAA vessels of TgSwDI mice in an unbiased manner (Supplementary File 2). All image analysis was performed by averaging the images from three tissue sections per mouse to represent each individual mouse.

#### Prussian blue staining

We used iron stain kit (abcam, ab150674) which detects ferric iron in tissues, formed after the lysis of RBCs. Ferric iron reacts with acid ferrocyanide producing a blue color (Prussian Blue stain reaction) (34). Prussian blue iron staining was performed as described previously (62, 63). Briefly, tissue sections from 18-months-old TgSwDI mice and WT littermates were fixed in 4% PFA in PBS for 20 minutes at RT. The sections were then incubated for 3 minutes in iron stain, which is a 1:1 mixture of potassium ferrocyanide and hydrochloric acid. The tissues were washed thoroughly with distilled water, followed by counterstaining with nuclear fast red solution for 5 minutes. The sections were then washed again 4 times with distilled water, 5 minutes each and then dehydrated in 95% ethanol followed by absolute alcohol. The sections were then mounted with VectaMount™ Mounting Medium (Vectorlabs, H-5000) followed by visualization under the brightfield microscope.

#### Evans blue assay

Blood brain barrier integrity was observed using Evans blue dye as described previously with slight modification (64). A 2% solution of Evans Blue dye in normal saline was injected intraperitoneally in 18-months-old TgSwDI and WT littermates (10ul dye/g of body weight). The dye was allowed to circulate for 6 hours. Afterwards, the mice were transcardially perfused with 30ml of ice-cold saline, plus 2.5% heparin solution and the brain tissue were removed, weighed, homogenized in 50% TCA in 1:3 weight to volume ratio and incubated for 1 hour on ice.

Samples were centrifuged for 30 minutes at 10000g at 4° C. 30ul of supernatant and 90ul of 95% ethanol was added in single well of 96 well plate. Each sample was in triplicate. Evans Blue dye content in brain tissue was measured by reading absorbance at 620 nm and quantified according to a standard curve. The results are presented as ng of Evans Blue stain/mg of tissue.

#### Barnes Maze Test

Spatial memory was measured using a Barnes maze as previously described (48) with slight modifications. The test was performed on 12 months-old siRNA-LNP treated mice. Testing room contained stationary visual cues, and the tester remained in a specific position during testing. The maze consisted of a white circular platform 92 cm in diameter with 20 holes (5.5 cm diameter) evenly spaced around the perimeter. One hole was assigned as a target and led to a dark escape box. All other holes had false bottoms. For motivation, the maze was illuminated with two halogen lamps in addition to room lighting producing an illumination of 1200 lux at the maze surface. The mice were placed at the center of the platform and given 300 seconds to explore and locate the target hole. If a mouse failed to enter the target hole within this time, it was gently guided to the escape box, allowed to enter, and kept there for 30 seconds before being returned to its home cage. If the mouse entered the target hole, it was immediately returned to its home cage. The mice underwent two trials per day for three consecutive days using the same training method. Following this, probe trials were conducted 24 hours after the final training session. During the probe trials, all the holes were closed, and the mice were allowed to explore for 120 seconds. The mouse’s movements were monitored with a camera interfaced to a computer which operated video tracking software (Noldus, EthoVision XT 14). An examination of the latency to find the target hole and cumulative duration in target hole was analyzed.

#### Western blot Assay

Brain tissue form 18-months-old TgSwDI mice and WT littermates, and 12-months-old siRNA treated TgSwDI mice and WT littermates were lysed in radioimmunoprecipitation assay (RIPA) buffer (25 mM Tris•HCl pH 7.6, 150 mM NaCl, 1% NP-40, 1% sodium deoxycholate, 0.1% SDS (Thermofisher scientific, #89900) along with protease and phosphatase inhibitor cocktail (Thermofisher #A32962) and centrifugated for 15 min at 10,000 rpm, 4°C. Tissue lysates were quantified using the BCA protein assay (Pierce #23223) and samples for western blot was prepared. 30 µg of protein was loaded per well on Bolt™ Bis-Tris Plus Mini Protein Gels, 4-12% (Thermofisher scientific, #NW04125BOX) with Bolt MES SDS Running Buffer (Invitrogen, #B0002). Gels were transferred onto Immobilon®-P PVDF Membrane (Millipore sigma, #IPVH85R) using Tris transfer buffer [400 mmol/mL Tris base, 70 mmol/mL glycine, 10% methanol]. Membrane was blocked in 5% milk in TBST (Tris Buffer salin+0.1% Tween-20) at RT for 1 hour on orbital shaker. Thereafter, membranes were incubated with primary antibodies diluted in 5% milk TBST (Tris Buffer salin+0.1% Tween-20) overnight at 4°C on shaker. Primary antibodies used were rabbit anti-ZO-1 (1∶1000, Invitrogen #40-2200), rabbit anti-Claudin-5 (1/1000, Invitrogen # 341600), rabbit anti-Occludin (1/1000 Abcam # ab167161), rabbit anti-fibrinogen (1/5000, Dako, A008002), mouse anti GAPDH (1/10000, Abcam, ab9484) and rabbit anti-transferrin (1/1000, Cell signalling # 35293). Next, membranes were washed with TBST (Tris Buffer salin+0.1% Tween-20), 3 times for 10 minutes each. Then, membranes were incubated with secondary antibodies: donkey anti rabbit IgG conjugated to horseradish peroxidase (HRP) (1/20000, Cytiva, NA934V) and sheep anti mouse IgG conjugated to HRP (Cytiva, NA931V) diluted in 5% milk TBST for 1 hour at RT on shaker. Membranes were then washed with TBST, 10 minutes on shaker and developed by using Pierce™ ECL Plus Western Blotting Substrate (Thermo fisher #32134). PVDF membranes were imaged using iBright™ CL1500 Imaging System and analyzed by densitometry using Fiji-Image J software. Data was expressed in arbitrary densitometry units (ADU).

## Statistical analysis

Statistical analysis was performed using GraphPad Prism 8 (GraphPad Software, San Diego, CA). Statistical analyses were conducted using Prism 8. All numerical values presented in graphs are mean ± SEM. Statistical analyses were assessed as described in the text using either one-way or two-way ANOVA followed by Bonferroni multiple comparisons post-hoc analysis or two-tailed unpaired t test. P values below 0.05 were considered significant. P values were as follows: ***P < 0.001; **P < 0.01; *P < 0.05.

## Acknowledgments

This work was supported by NIH Grants NS104386 (to H.J.A.), AG078245 (to H.J.A.), and NIH grant HL168009 (to C.J.K.). We thank the Cellular Imaging and Histology Core, Rutgers, Newark for assistance with confocal microscopy. We also thank Kelly Mulraney, veterinary staff at In Vivo Research Services (IVRS) core, Rutgers, for assisting with tail vein injections of siRNA LNP’s. We thank members of the Ahn laboratory for scientific discussion, and experimental support.

## Authorship Contribution

V.S., C.J.K., and H.J.A. designed research; V.S., N.R., K. G., F.F., and A.C., performed research; V.S., N.R., and H.J.A. analyzed data; and V.S., A.C., and H.J.A. wrote the paper.

## Declaration of Interests

C.J.K., is a director, shareholder and co-founder of companies developing RNA-therapies, SeraGene Therapeutics, Inc. and NanoVation Therapeutics, Inc. C.J.K and F.F. have filed intellectual property on RNA-based therapies with the intention of commercializing these inventions.

**Correspondence:** Hyung Jin Ahn, Department of Pharmacology, Physiology and Neurosciences, Rutgers-New Jersey Medical School, Newark, NJ, USA;

## References

1. C. C. Hays, Z. Z. Zlatar, C. E. Wierenga, The Utility of Cerebral Blood Flow as a Biomarker of Preclinical Alzheimer’s Disease. Cell Mol Neurobiol 36, 167–179 (2016).

2. M. A. Binnewijzend et al., Cerebral perfusion in the predementia stages of Alzheimer’s disease. Eur Radiol 26, 506–514 (2016).

3. R. D. Bell, B. V. Zlokovic, Neurovascular mechanisms and blood-brain barrier disorder in Alzheimer’s disease. Acta Neuropathol 118, 103–113 (2009).

4. N. A. Johnson et al., Pattern of cerebral hypoperfusion in Alzheimer disease and mild cognitive impairment measured with arterial spin-labeling MR imaging: initial experience. Radiology 234, 851–859 (2005).

5. A. Serrano-Pozo et al., Examination of the clinicopathologic continuum of Alzheimer disease in the autopsy cohort of the National Alzheimer Coordinating Center. J Neuropathol Exp Neurol 72, 1182–1192 (2013).

6. M. Yamada, Cerebral amyloid angiopathy: emerging concepts. J Stroke 17, 17–30 (2015).

7. D. R. Thal et al., Capillary cerebral amyloid angiopathy is associated with vessel occlusion and cerebral blood flow disturbances. Neurobiol Aging 30, 1936–1948 (2009).

8. E. E. Smith et al., Impaired visual evoked flow velocity response in cerebral amyloid angiopathy. Neurology 71, 1424–1430 (2008).

9. J. P. Vonsattel et al., Cerebral amyloid angiopathy without and with cerebral hemorrhages: a comparative histological study. Ann Neurol 30, 637–649 (1991).

10. A. Lauer et al., Microbleeds on MRI are associated with microinfarcts on autopsy in cerebral amyloid angiopathy. Neurology 87, 1488–1492 (2016).

11. A. Viswanathan, S. M. Greenberg, Cerebral amyloid angiopathy in the elderly. Ann Neurol 70, 871–880 (2011).

12. K. Gouveia-Freitas, A. J. Bastos-Leite, Perivascular spaces and brain waste clearance systems: relevance for neurodegenerative and cerebrovascular pathology. Neuroradiology 63, 1581–1597 (2021).

13. N. J. Abbott, M. E. Pizzo, J. E. Preston, D. Janigro, R. G. Thorne, The role of brain barriers in fluid movement in the CNS: is there a ’glymphatic’ system? Acta Neuropathol 135, 387–407 (2018).

14. M. K. Rasmussen, H. Mestre, M. Nedergaard, The glymphatic pathway in neurological disorders. Lancet Neurol 17, 1016–1024 (2018).

15. S. Peng, J. Liu, C. Liang, L. Yang, G. Wang, Aquaporin-4 in glymphatic system, and its implication for central nervous system disorders. Neurobiol Dis 179, 106035 (2023).

16. R. Owasil et al., The Pattern of AQP4 Expression in the Ageing Human Brain and in Cerebral Amyloid Angiopathy. Int J Mol Sci 21 (2020).

17. M. D. Manescu et al., Aquaporin 4 modulation drives amyloid burden and cognitive abilities in an APPPS1 mouse model of Alzheimer’s disease. Alzheimers Dement 21, e70164 (2025).

18. P. Marazuela et al., Circulating AQP4 Levels in Patients with Cerebral Amyloid Angiopathy-Associated Intracerebral Hemorrhage. J Clin Med 10 (2021).

19. Y. L. Lan, J. Zhao, T. Ma, S. Li, The Potential Roles of Aquaporin 4 in Alzheimer’s Disease. Mol Neurobiol 53, 5300–5309 (2016).

20. C. Yang et al., Aquaporin-4 and Alzheimer’s Disease. J Alzheimers Dis 52, 391–402 (2016).

21. D. M. Wilcock, M. P. Vitek, C. A. Colton, Vascular amyloid alters astrocytic water and potassium channels in mouse models and humans with Alzheimer’s disease. Neuroscience 159, 1055–1069 (2009).

22. M. Simon et al., Loss of perivascular aquaporin-4 localization impairs glymphatic exchange and promotes amyloid beta plaque formation in mice. Alzheimers Res Ther 14, 59 (2022).

23. M. Cortes-Canteli et al., Fibrinogen and beta-amyloid association alters thrombosis and fibrinolysis: a possible contributing factor to Alzheimer’s disease. Neuron 66, 695–709 (2010).

24. H. J. Ahn, S. K. Baker, E. H. Norris, S. Strickland, Inflaming the Brain. Neuron 101, 991–993 (2019).

25. J. Paul, S. Strickland, J. P. Melchor, Fibrin deposition accelerates neurovascular damage and neuroinflammation in mouse models of Alzheimer’s disease. J Exp Med 204, 1999–2008 (2007).

26. M. Cortes-Canteli, L. Mattei, A. T. Richards, E. H. Norris, S. Strickland, Fibrin deposited in the Alzheimer’s disease brain promotes neuronal degeneration. Neurobiol Aging 36, 608–617 (2015).

27. S. A. Cajamarca, E. H. Norris, L. van der Weerd, S. Strickland, H. J. Ahn, Cerebral amyloid angiopathy-linked beta-amyloid mutations promote cerebral fibrin deposits via increased binding affinity for fibrinogen. Proc Natl Acad Sci U S A 117, 14482–14492 (2020).

28. P. K. Eide, H. A. Hansson, Blood-brain barrier leakage of blood proteins in idiopathic normal pressure hydrocephalus. Brain Res 1727, 146547 (2020).

29. A. C. Yang et al., Physiological blood-brain transport is impaired with age by a shift in transcytosis. Nature 583, 425–430 (2020).

30. P. G. Frank, S. Pavlides, M. P. Lisanti, Caveolae and transcytosis in endothelial cells: role in atherosclerosis. Cell Tissue Res 335, 41–47 (2009).

31. N. Muradashvili, R. Tyagi, N. Tyagi, S. C. Tyagi, D. Lominadze, Cerebrovascular disorders caused by hyperfibrinogenaemia. J Physiol 594, 5941–5957 (2016).

32. N. Muradashvili, R. L. Benton, R. Tyagi, S. C. Tyagi, D. Lominadze, Elevated level of fibrinogen increases caveolae formation; role of matrix metalloproteinase-9. Cell Biochem Biophys 69, 283–294 (2014).

33. J. Davis et al., Early-onset and robust cerebral microvascular accumulation of amyloid beta-protein in transgenic mice expressing low levels of a vasculotropic Dutch/Iowa mutant form of amyloid beta-protein precursor. J Biol Chem 279, 20296–20306 (2004).

34. S. Liu et al., Comparative analysis of H&E and Prussian blue staining in a mouse model of cerebral microbleeds. J Histochem Cytochem 62, 767–773 (2014).

35. Y. Okamoto et al., Cerebral hypoperfusion accelerates cerebral amyloid angiopathy and promotes cortical microinfarcts. Acta Neuropathol 123, 381–394 (2012).

36. S. Saito et al., Taxifolin inhibits amyloid-beta oligomer formation and fully restores vascular integrity and memory in cerebral amyloid angiopathy. Acta Neuropathol Commun 5, 26 (2017).

37. M. Wolman et al., Evaluation of the dye-protein tracers in pathophysiology of the blood-brain barrier. Acta Neuropathol 54, 55–61 (1981).

38. H. L. Wang, T. W. Lai, Optimization of Evans blue quantitation in limited rat tissue samples. Sci Rep 4, 6588 (2014).

39. P. C. Nahirney, P. Reeson, C. E. Brown, Ultrastructural analysis of blood-brain barrier breakdown in the peri-infarct zone in young adult and aged mice. J Cereb Blood Flow Metab 36, 413–425 (2016).

40. M. J. Haley, C. B. Lawrence, The blood-brain barrier after stroke: Structural studies and the role of transcytotic vesicles. J Cereb Blood Flow Metab 37, 456–470 (2017).

41. D. E. Levy et al., Ancrod in acute ischemic stroke: results of 500 subjects beginning treatment within 6 hours of stroke onset in the ancrod stroke program. Stroke 40, 3796–3803 (2009).

42. L. J. Juang et al., Suppression of fibrin(ogen)-driven pathologies in disease models through controlled knockdown by lipid nanoparticle delivery of siRNA. Blood 139, 1302–1311 (2022).

43. G. E. Hernandez et al., Aortic intimal resident macrophages are essential for maintenance of the non-thrombogenic intravascular state. Nat Cardiovasc Res 1, 67–84 (2022).

44. W. S. Hur et al., Elimination of fibrin polymer formation or crosslinking, but not fibrinogen deficiency, is protective against diet-induced obesity and associated pathologies. J Thromb Haemost 20, 2873–2886 (2022).

45. W. S. Hur et al., Hypofibrinogenemia with preserved hemostasis and protection from thrombosis in mice with an Fga truncation mutation. Blood 139, 1374–1388 (2022).

46. F. Xu et al., Early-onset subicular microvascular amyloid and neuroinflammation correlate with behavioral deficits in vasculotropic mutant amyloid beta-protein precursor transgenic mice. Neuroscience 146, 98–107 (2007).

47. L. S. Robison et al., Long-term voluntary wheel running does not alter vascular amyloid burden but reduces neuroinflammation in the Tg-SwDI mouse model of cerebral amyloid angiopathy. J Neuroinflammation 16, 144 (2019).

48. S. E. Setti, T. Flanigan, J. Hanig, S. Sarkar, Assessment of sex-related neuropathology and cognitive deficits in the Tg-SwDI mouse model of Alzheimer’s disease. Behav Brain Res 428, 113882 (2022).

49. S. Rotshenker, Microglia and macrophage activation and the regulation of complement-receptor-3 (CR3/MAC-1)-mediated myelin phagocytosis in injury and disease. J Mol Neurosci 21, 65–72 (2003).

50. D. Davalos et al., Fibrinogen-induced perivascular microglial clustering is required for the development of axonal damage in neuroinflammation. Nat Commun 3, 1227 (2012).

51. J. Miao et al., Cerebral microvascular amyloid beta protein deposition induces vascular degeneration and neuroinflammation in transgenic mice expressing human vasculotropic mutant amyloid beta precursor protein. Am J Pathol 167, 505–515 (2005).

52. B. J. Andreone et al., Blood-Brain Barrier Permeability Is Regulated by Lipid Transport-Dependent Suppression of Caveolae-Mediated Transcytosis. Neuron 94, 581–594 e585 (2017).

53. B. W. Chow, C. Gu, Gradual Suppression of Transcytosis Governs Functional Blood-Retinal Barrier Formation. Neuron 93, 1325–1333 e1323 (2017).

54. S. B. Gaudreault, D. Dea, J. Poirier, Increased caveolin-1 expression in Alzheimer’s disease brain. Neurobiol Aging 25, 753–759 (2004).

55. T. Y. Ha et al., Age-related increase in caveolin-1 expression facilitates cell-to-cell transmission of alpha-synuclein in neurons. Mol Brain 14, 122 (2021).

56. M. J. Kang et al., Caveolin-1 upregulation in senescent neurons alters amyloid precursor protein processing. Exp Mol Med 38, 126–133 (2006).

57. J. J. Iliff et al., Impairment of glymphatic pathway function promotes tau pathology after traumatic brain injury. J Neurosci 34, 16180–16193 (2014).

58. K. Ishida et al., Glymphatic system clears extracellular tau and protects from tau aggregation and neurodegeneration. J Exp Med 219 (2022).

59. T. J. Pedersen, S. A. Keil, W. Han, M. X. Wang, J. J. Iliff, The effect of aquaporin-4 mis-localization on Abeta deposition in mice. Neurobiol Dis 181, 106100 (2023).

60. B. C. Reeves et al., Glymphatic System Impairment in Alzheimer’s Disease and Idiopathic Normal Pressure Hydrocephalus. Trends Mol Med 26, 285–295 (2020).

61. F. Ferraresso et al., Comparison of DLin-MC3-DMA and ALC-0315 for siRNA Delivery to Hepatocytes and Hepatic Stellate Cells. Mol Pharm 19, 2175–2182 (2022).

62. G. Gomori, Microtechnical Demonstration of Iron: A Criticism of its Methods. Am J Pathol 12, 655–664 651 (1936).

63. D. T. Winkler et al., Spontaneous hemorrhagic stroke in a mouse model of cerebral amyloid angiopathy. J Neurosci 21, 1619–1627 (2001).

64. A. Manaenko, H. Chen, J. Kammer, J. H. Zhang, J. Tang, Comparison Evans Blue injection routes: Intravenous versus intraperitoneal, for measurement of blood-brain barrier in a mice hemorrhage model. J Neurosci Methods 195, 206–210 (2011).

