## Supplementary Figures for "Aquaporin-4 and Caveolin-1 as Mediators of Fibrinogen-Driven Cerebrovascular Pathology in Cerebral Amyloid Angiopathy"

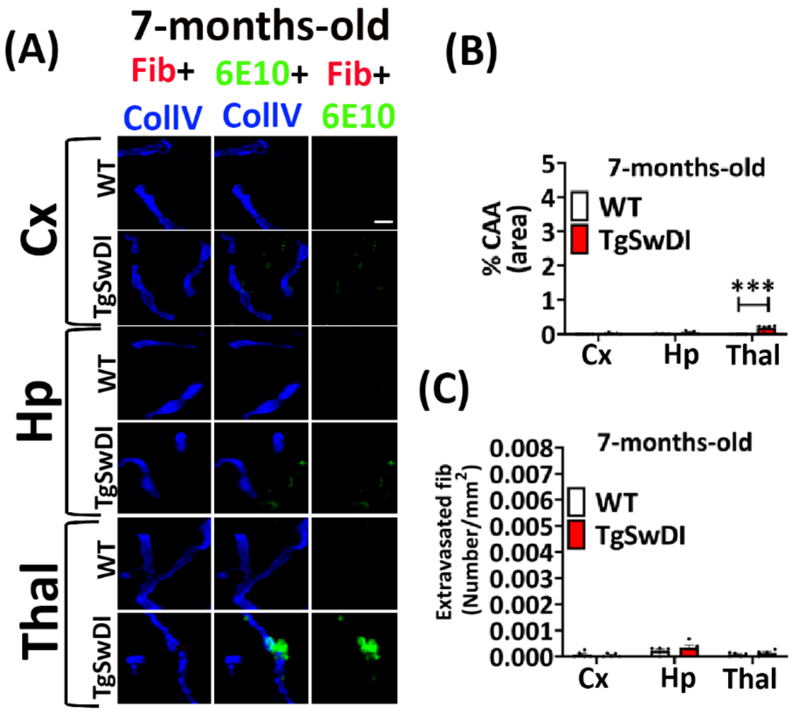


Figure S1. Increased CAA in the Thalamus of 7-month-old TgSwDI mice. (A) Representative images of the brain of 7-months-old TgSwDI and WT littermates probed with antibodies against fibrin(ogen) (red), Aβ (6E10, green), and blood vessels (collagen IV (CollV), blue). CAA is represented as colocalized collagen IV and Aβ. Statistical quantification of CAA (B) and the number of extravasated fibrin(ogen) (C) as observed in (A). (n=5 mice per group). Data were analyzed by using two-way ANOVA with the Bonferroni post hoc test and shown as average ± SEM; ***P < 0.001.

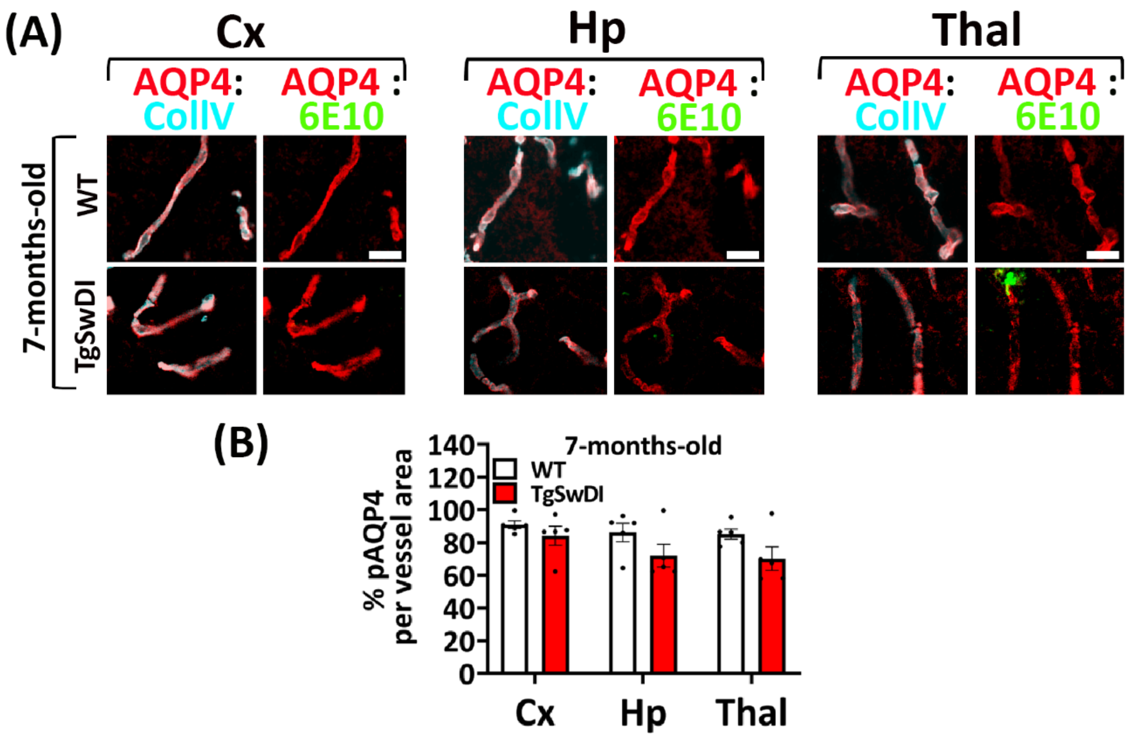


Figure S2. No significant change in perivascular AQP4 levels in the brains of 7-month-old TgSwDI mice. (A) Representative images were stained with antibodies against AQP4 (red), Aβ (6E10, green), and blood vessels (collagen IV (CollV), cyan). Perivascular AQP4 is visualized as pink. Scale=10μm. (B) Statistical quantification of perivascular AQP4 in 7-months-old TgSwDI and WT littermates. (n=5 mice per group). No significant loss of perivascular AQP4 was observed in TgSwDI mice compared to WT littermates. Data were analyzed by using two-way ANOVA with the Bonferroni post hoc test and shown as average ± SEM.

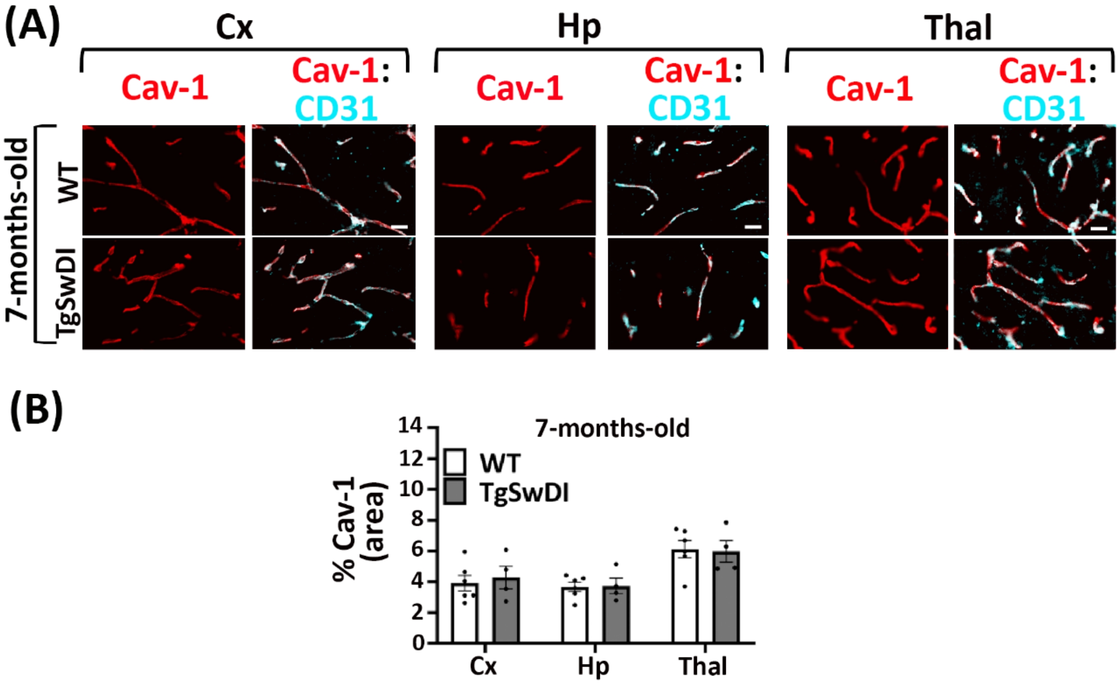


Figure S3. Non-significant change in Cav-1 levels observed in the brain of 7-months-old TgSwDI mice. (A) Representative images stained with antibodies against Cav-1 (red) and endothelial cell marker, CD31 (cyan). Scale=20μm. (B) Statistical quantification of Cav-1, as shown in (A), revealed no significant difference in Cav-1 levels between TgSwDI mice and WT littermates at 7 months of age (n=4-6 mice per group). Data were analyzed by using two-way ANOVA with the Bonferroni post hoc test and shown as average ± SEM.

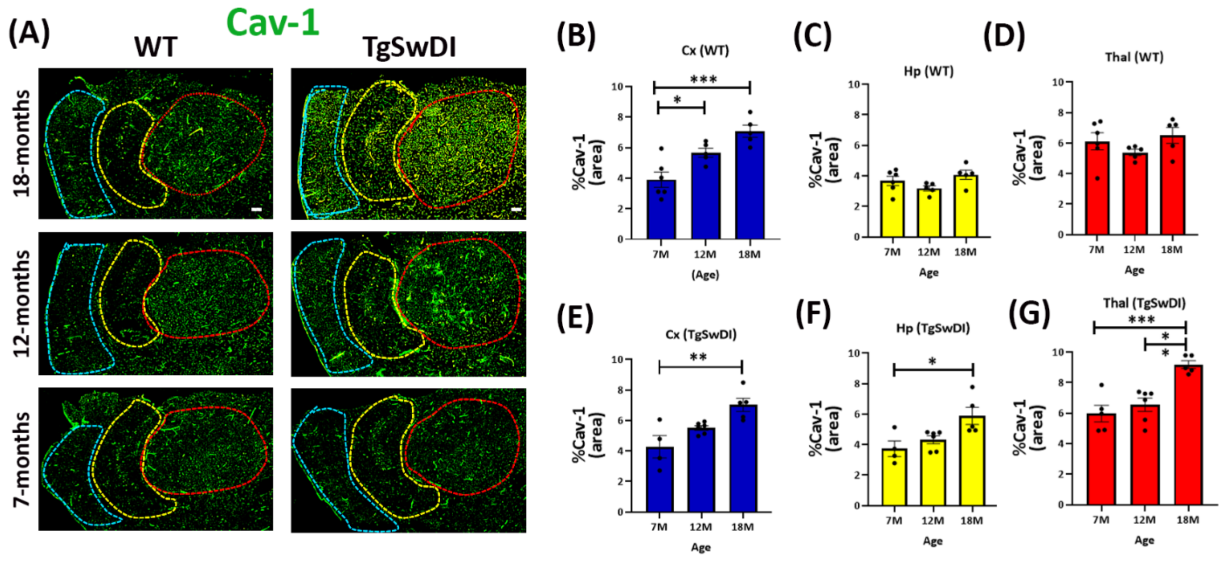


Figure S4. Age-dependent increase of Cav-1 in TgSwDI mice. (A) Representative images of brain coronal sections from 7-, 12-, and 18-month-old TgSwDI and WT littermates, immunostained with antibodies against Cav-1. The cortex, hippocampus, and thalamus are outlined with blue, yellow, and red boundaries, respectively. (B-G) Statistical quantification of Cav-1, as shown in (A). (n=4-5 mice per group). Scale=400μm. Data were analyzed by using two-way ANOVA with the Bonferroni post hoc test and shown as average ± SEM; *, $P < 0.05$, **, $P < 0.01$, ***$P < 0.001$.

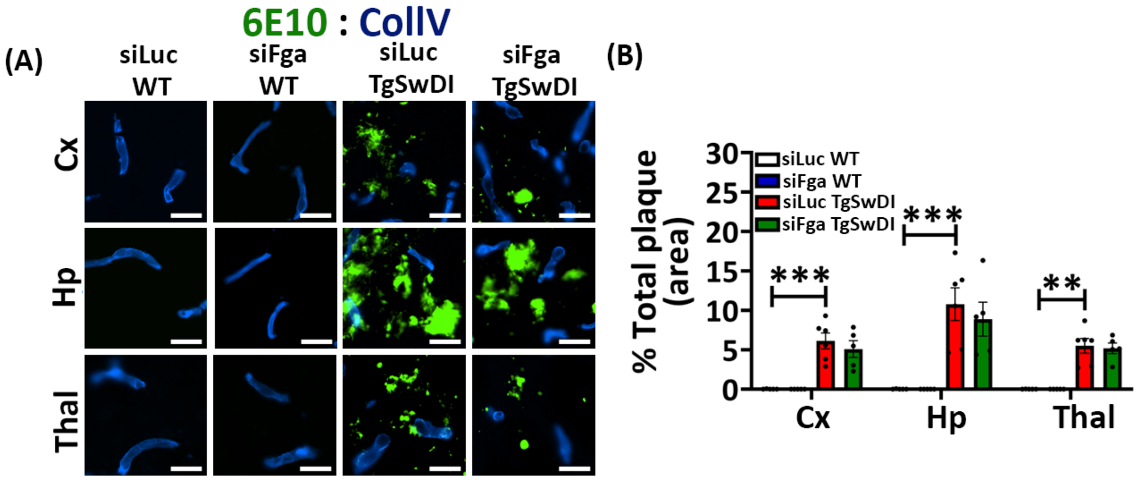


Figure S5. No significant change of amyloid plaque levels in fibrinogen depleted TgSwDI mice. (A) Coronal brain sections from siRNA-treated mice were immunostained for amyloid plaques (6E10, green) and blood vessels (Collagen IV (CollV), blue). (B) Statistical quantification showed that siLuc TgSwDI mice exhibited significantly higher amyloid plaques as compared to siLuc WT, with no significant difference in amyloid plaque levels between siFga TgSwDI and siLuc TgSwDI mice (n=5-6 mice per group). Scale=20μm. Data were analyzed by using two-way ANOVA with the Bonferroni post hoc test and shown as average ± SEM; **, P < 0.01, ***P < 0.001.

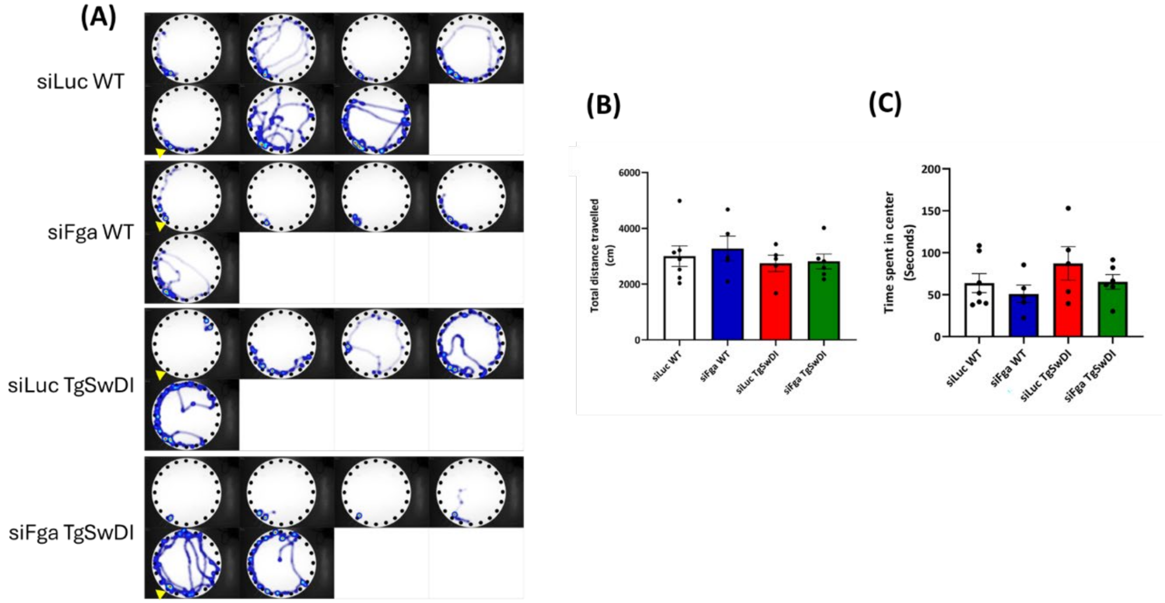


Figure S6. Fibrinogen-depleted TgSwDI mice exhibited impaired spatial memory but showed no deficits in locomotion or anxiety-related behaviors. (A) Barnes maze heatmaps for each mouse (one maze per mouse) depict movement paths and time spent in different areas using a color gradient. Warmer colors (red/yellow) indicate regions where mice spent more time, while cooler colors (blue) represent areas with less time spent. Groups include siLuc WT (n=7), siFga WT (n=5), siLuc TgSwDI (n= 5), and siFga TgSwDI (n=6). Yellow arrowhead indicates the target hole in the maze. (B & C) Open field task showed that there was no significant difference in total distance travelled (B) and time spent in center (measure of anxiety) (C) between any group (n=5-7).

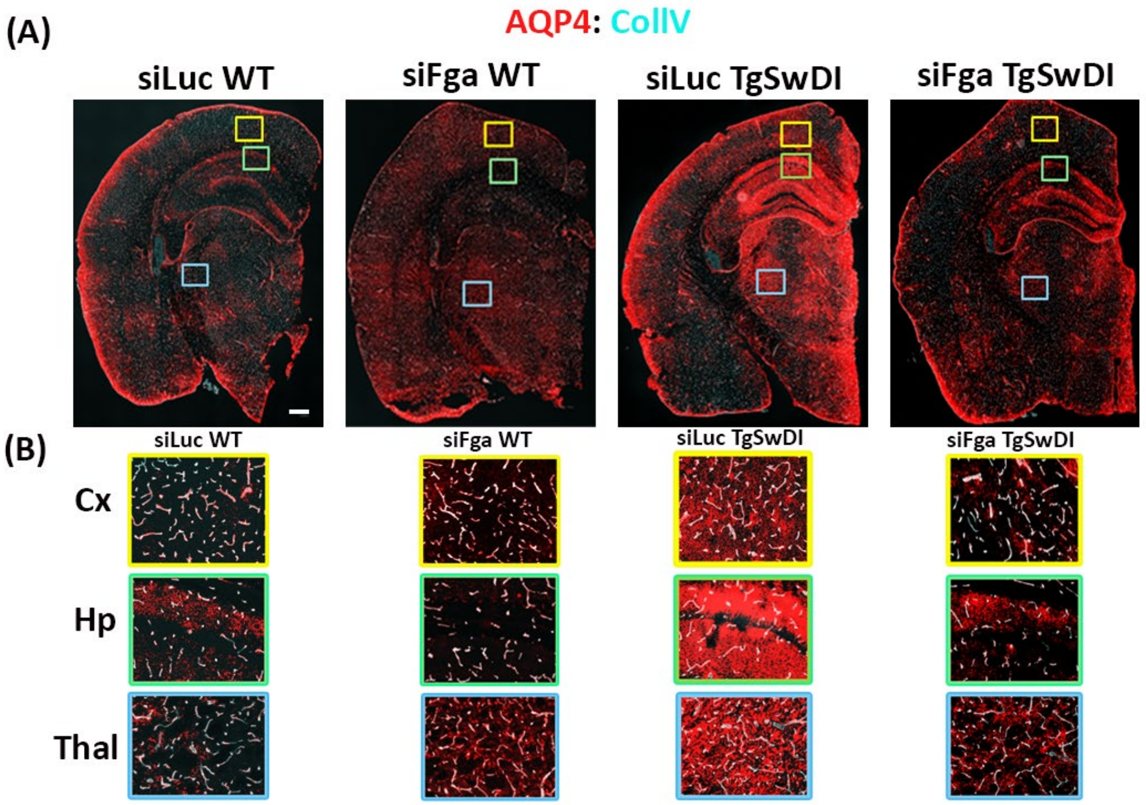


Figure S7. Fibrinogen depletion reduces the elevated levels of non-vascular AQP4 in TgSwDI mice. (A) Representative images of brain coronal sections of TgSwDI and WT littermates treated with siLuc and siFga LNP's and probed with antibodies against AQP4 (red) plus blood vessels (collagen IV, cyan). (n=5-7 per group). Scale=400μm. (B) The yellow box shows a zoom-in image of the cortex, the green box highlights the hippocampus, and the blue box indicates the thalamus.

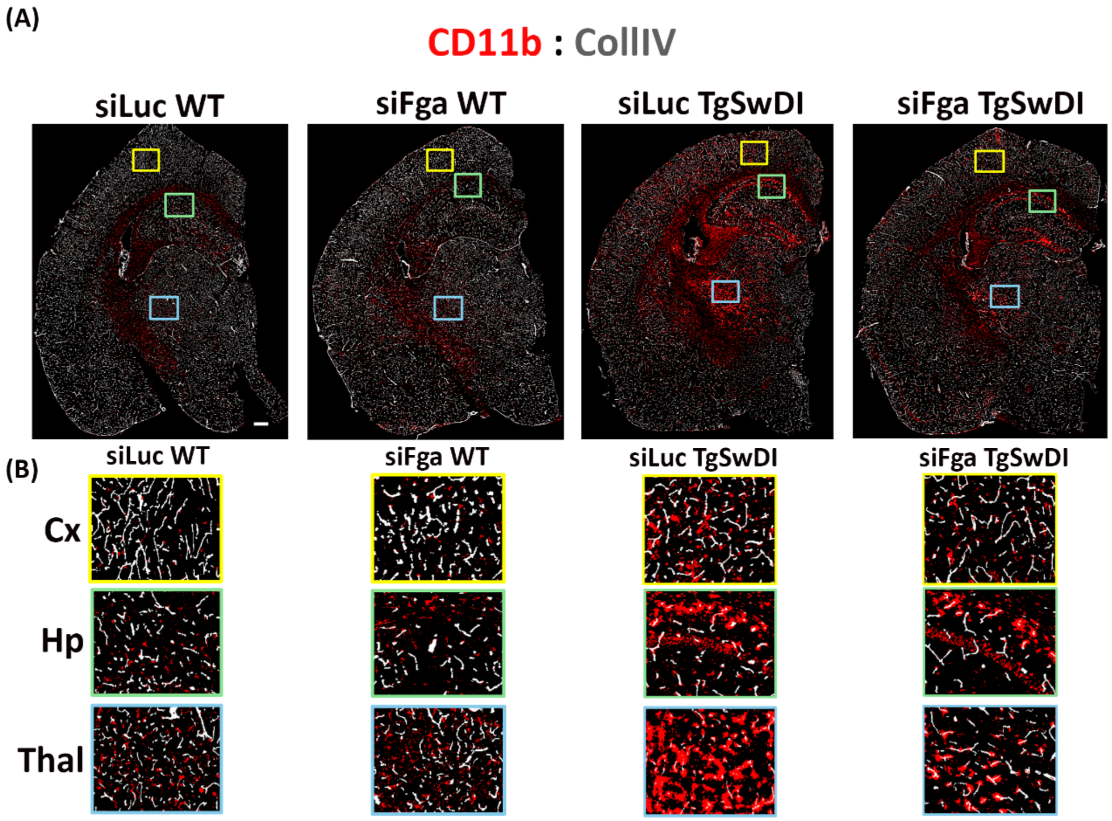


Figure S8. Fibrinogen depletion reduces the activated microglial cells (CD11b positive) in TgSwDI mice. (A) Representative images of brain coronal sections of TgSwDI and WT littermates treated with siLuc and siFga LNP's and probed with antibodies against CD11b (red) plus blood vessels (collagen IV, grey). (n=5-7 per group) Scale=400μm. (B) The yellow box shows a zoom-in image of the cortex, the green box highlights the hippocampus, and the blue box indicates the thalamus.

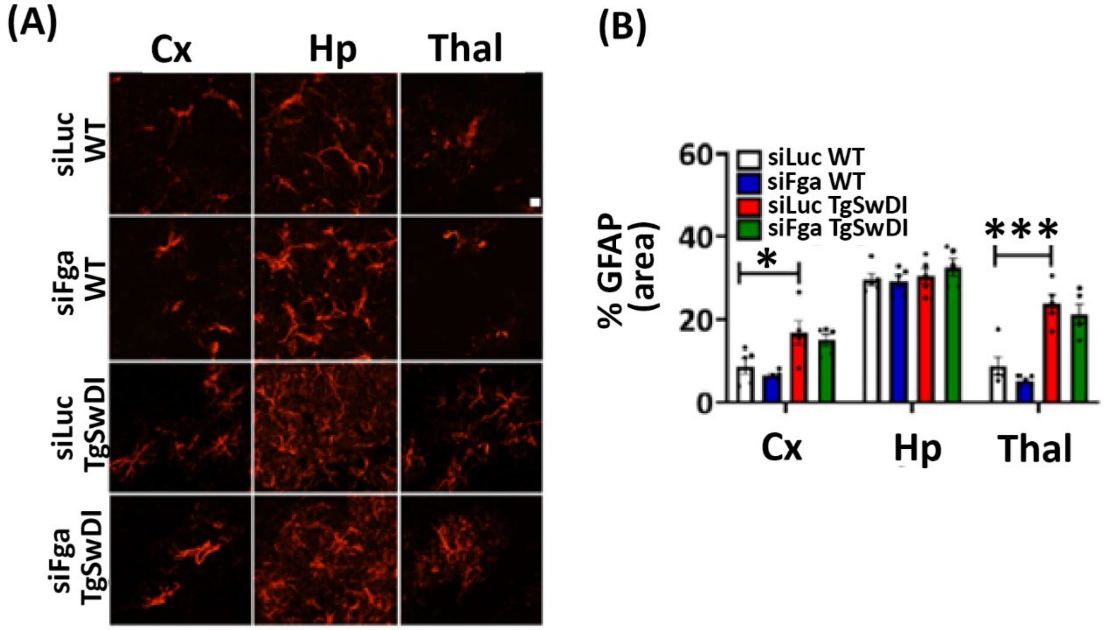


Figure S9. No significant change in reactive astrocytes was observed in fibrinogen-depleted TgSwDI mice. (A) Coronal brain sections from siRNA-treated mice were immunostained for GFAP to assess reactive astrocyte levels. (B) Statistical quantification of (A) shows that siLuc TgSwDI mice exhibited significantly higher GFAP expression in the cortex and thalamus compared to siLuc WT mice, but no significant differences were observed between siFga TgSwDI and siLuc TgSwDI mice. (n=5 mice per group). Scale=10 μm. Data were analyzed by using two-way ANOVA with the Bonferroni post hoc test and shown as average ± SEM; *, $P < 0.05$, ***$P < 0.001$.

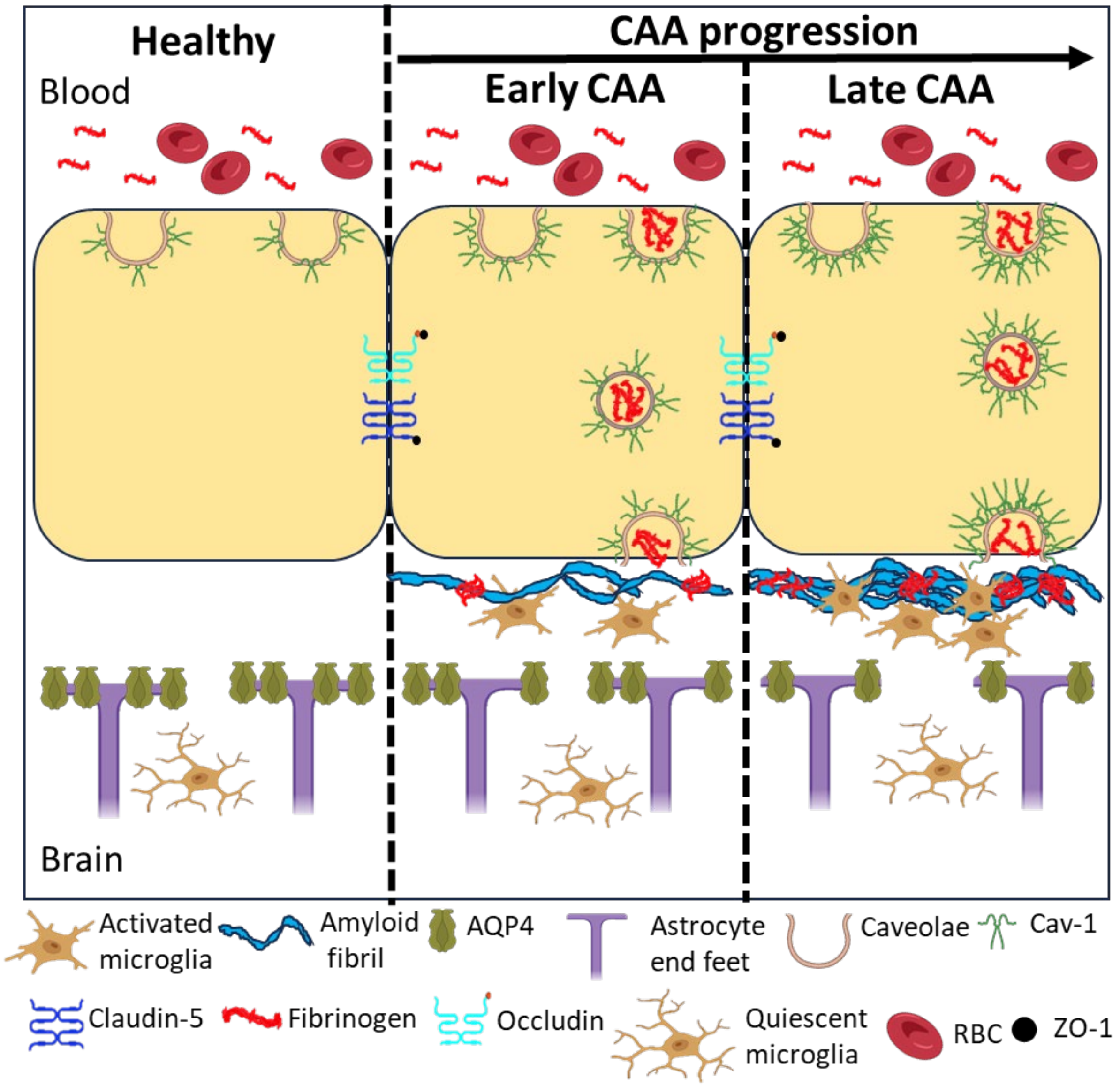


Figure S10. Schematic illustration of fibrinogen-driven cerebrovascular pathology in cerebral amyloid angiopathy. Under normal conditions, the BBB remains intact, characterized by stable tight junction proteins and physiological levels of Cav-1 in endothelial cells. No fibrinogen extravasation, CAA, or microglial activation is observed. Aquaporin-4 (AQP4) is properly polarized to astrocyte endfeet in the PVS. As CAA progresses and with aging, endothelial cells show elevated Cav-1 expression, which may facilitate fibrinogen extravasation through Cav-1-coated caveolae vesicles, potentially colocalizing with Aβ in the PVS. In response, perivascular microglia become activated, adopting a pro-inflammatory phenotype, while AQP4 polarization is disrupted, impairing glymphatic clearance. In advanced CAA and aging, cumulative vascular damage leads to widespread Aβ deposition, chronic fibrinogen leakage, and persistent neuroinflammation. These changes contribute to prolonged microglial activation, perivascular gliosis and AQP4 depolarization, impairing homeostatic fluid dynamics and leading to reduced clearance of toxic proteins. The combined effects of aging and CAA ultimately promote vascular dysfunction, white matter damage, and neurodegenerative processes.
