## Supplemental Code for "Aquaporin-4 and Caveolin-1 as Mediators of Fibrinogen-Driven Cerebrovascular Pathology in Cerebral Amyloid Angiopathy": Matlab_code_cd11b.m

```
close all
results = Image_Crop("G:\cd11b_matlab_images_rdrive\8865_m_fib_wt", 'G', 'B',  160, 100, [ "cort", "hippo", "thal" ] , 0.1);
function [results] = Image_Crop(Directory,color_target,color_criteria, target_thresh, criteria_thresh, regions, top_thresh)
close all
if color_target == 'R'
    color_target = 1;
end
if color_target == 'G'
    color_target = 2;
end
if color_target == 'B'
    color_target = 3;
end

if color_criteria == 'R'
    color_criteria = 1;
end
if color_criteria == 'G'
    color_criteria = 2;
end
if color_criteria == 'B'
    color_criteria = 3;
end
percent = top_thresh*100;
percent = string(percent);
rootdir = Directory;
filelist = dir(fullfile(rootdir, '**\*.*'));  %get list of files and folders in any subfolder
filelist = filelist(~[filelist.isdir]);
filename = [];
for m = 1:length(filelist)
    hold = filelist(m).name;
    hold = string(hold);
    filename = [filename, hold];
    %print(filename);
end
% filter = contains(filename, "Composite"); %for normal
filter = contains(filename, "RGB"); %for others
filter_not = contains(filename, "microglia");
filter = logical(filter - filter_not);
filename = filename(filter);
filelist = filelist(filter);
filter = zeros(1,length(filename));
for i = 1:length(regions)
    region_filter = contains(filename, regions(i));
    filter = filter + region_filter;
end
filter = logical(filter);
filelist = filelist(filter);
results = struct('Name', {[]}, 'Target_area', {[]});
for m = 1:length(filelist)
    close all
    image_name = filelist(m).folder + "\" + filelist(m).name;
    t = Tiff(image_name, 'r');
    Image = read(t);
    figure
    imshow(Image)
    I_0 = Image;
    I_criteria = uint8(I_0(:,:,color_criteria) > criteria_thresh);
    I_target = uint8(I_0(:,:,color_target)>target_thresh);
 
    
    I_0(:,:,color_target) = I_target.*I_0(:,:,color_target);

    I_0(:,:,color_criteria) = I_criteria.*I_0(:,:,color_criteria);
    figure
    imshow(I_0)
    I_interest = I_0(:,:,color_target);

    bwthresh = 0;
    connection = 6;
    figure
    imshow(I_interest)

    I_criteria = uint8(I_0(:,:,color_criteria) > criteria_thresh);
    I_target = uint8(I_0(:,:,color_target)>target_thresh);

    I_interest = uint8(I_interest);

    CC = bwconncomp(I_interest, 8);
    S = regionprops(CC,'Area');
    L = labelmatrix(CC);
    I_interest = ismember(L,find([S.Area]>=50));
    figure
    imshow(I_interest)
    I_interest = uint8(I_interest);
    I_check = I_interest.*255;
    I_check_im = Image;
    I_check_im(:,:,2) = I_check_im(:,:,2) + I_check;
    figure
    imshow(I_check_im)
   
    boundaries = bwboundaries(I_interest);
    BW = im2bw(I_interest,0);
    figure
    imshow(BW)
  
    st = regionprops(BW, 'BoundingBox');
    xmin = zeros(length(st),1);
    xmax = zeros(length(st),1);
    ymin = zeros(length(st),1);
    ymax = zeros(length(st),1);
    sizes = size(I_interest);
    mask = zeros(sizes(1), sizes(2));
    mask_between = zeros(sizes(1),sizes(2));
    area_check = zeros(4,length(st));
    for i = 1:length(st)
        rectangle('Position',[st(i).BoundingBox(1),st(i).BoundingBox(2),st(i).BoundingBox(3),st(i).BoundingBox(4)], 'EdgeColor','r','LineWidth',2 )
        xmin(i) = floor(st(i).BoundingBox(1));
        xmax(i) = xmin(i) + st(i).BoundingBox(3);
        ymin(i) = floor(st(i).BoundingBox(2));
        ymax(i) = ymin(i) + st(i).BoundingBox(4);
    end
    zero_check = xmin == 0;
    xmin = xmin + zero_check;
    zero_check = ymin == 0;
    ymin = ymin + zero_check;
    for i = 1:length(st)
        i
        object = boundaries{i,1};
        potential_mask = zeros(sizes(1),sizes(2));
        potential_mask = gpuArray(potential_mask);
        for j = 1:length(object)
            potential_mask(object(j,1),object(j,2)) = 1;
        end
        potential_mask = imfill(potential_mask);
        object = uint8(potential_mask);
        object = gpuArray(object);%gpu
        
        object = object.*I_0;
        object_mask = object(:,:,color_target)>target_thresh;
        object_mask = uint8(object_mask);
        target_mask = object(:,:,color_criteria)>criteria_thresh;
        target_mask = uint8(target_mask);
        mask_temp = zeros(sizes(1),sizes(2));
        mask_between_temp = zeros(sizes(1),sizes(2));
        area_check(1,i) = sum(sum(target_mask));
        area_check(2,i) = sum(sum(object_mask));
        area_check(3,i) = area_check(1,i)/area_check(2,i);
        area_check(4,i) = (ymax(i)-ymin(i))*(xmax(i)-xmin(i));
        if(sum(sum(target_mask)) >= sum(sum(object_mask))*top_thresh)
            mask_temp = potential_mask;

        end
        mask = mask + mask_temp;
        
    end
    mask_CAA = mask >0;
    figure
    imshow(mask_CAA)

    mask_reversed = mask == 0;

    I_interest = uint8(I_interest);
    
    CAA_mask = I_interest.*uint8(mask_CAA);
    CAA_mask = uint8(CAA_mask);
    CAA_Image = Image.*CAA_mask;
    area = sum(sum(mask_CAA))/2.74;
    I_check = uint8(CAA_mask).*255;
    I_check_im = Image;
    I_check_im(:,:,2) = I_check_im(:,:,2) + I_check;
    figure
    imshow(I_check_im)
    ind = strfind(filelist(m).name,"RGB");
    imname = char(filelist(m).folder);
    indexing = strfind(filelist(m).folder, "\");
    imname = string(imname(indexing(end)+1:end)) + filelist(m).name;
    imname = char(imname);
    folder_name = imname(1:ind-2);
    folder_name = filelist(m).folder + "\" + folder_name;
    if ~isfolder(folder_name)
        mkdir(folder_name)
    end
    image_name = folder_name + imname;
    image_name = char(image_name);
    results(end+1) = struct('Name', {imname}, 'Target_area', {area});
    imwrite(I_check_im, image_name(1:end-4)+percent+"microglia_area.tif",'tiff');

end
end
```
